# Functional and comparative genome analysis reveals clade-specific genome innovations in the killer fungus *Candida auris*

**DOI:** 10.1101/2021.01.29.428909

**Authors:** Aswathy Narayanan, Rakesh Netha Vadnala, Promit Ganguly, Pavitra Selvakumar, Shivaprakash M Rudramurthy, Rajendra Prasad, Arunaloke Chakrabarti, Rahul Siddharthan, Kaustuv Sanyal

**Affiliations:** Molecular Mycology Laboratory, Molecular Biology and Genetics Unit, Jawaharlal Nehru Centre for Advanced Scientific Research, Bangalore, India; Computational Biology, The Institute of Mathematical Sciences/HBNI, Chennai, India; Postgraduate Institute of Medical Education and Research, Chandigarh, India; Amity Institute of Biotechnology, Haryana, India

**Keywords:** *Candida haemulonii*, *Candida duobushaemulonii*, *Candida pseudohaemulonii*, *Candida lusitaniae*, *Candida fructus*, chromosome number reduction, centromere relocation, ancestral genome

## Abstract

The thermotolerant multidrug-resistant ascomycete *Candida auris* rapidly emerged since 2009 and simultaneously evolved in different geographical zones worldwide, causing superficial as well as systemic infections. The molecular events that orchestrated this sudden emergence of the killer fungus remain mostly elusive. Here, we identify centromeres in *C. auris* and related species, using a combined approach of chromatin immunoprecipitation and comparative genomic analyses. We find that *C. auris* and multiple other species in the *Clavispora/Candida* clade shared a conserved small regional centromere landscape lacking pericentromeres. Further, a centromere inactivation event led to karyotypic alterations in this species complex. Inter-species genome analysis identified several structural chromosomal changes around centromeres. In addition, centromeres are found to be rapidly evolving loci among the different geographical clades of the same species of *C. auris*. Finally, we reveal an evolutionary trajectory of the unique karyotype associated with clade 2 that consists of the drug susceptible isolates of *C. auris*.

## Introduction

First isolated from an infected ear of a patient in Japan in 2009*, Candida auris* emerged as a multidrug-resistant opportunistic fungal pathogen causing nosocomial infections worldwide in a short time span (Satoh et al. 2009; Schelenz et al. 2016; Morales-López et al. 2017; Vallabhaneni et al. 2017; Ruiz-Gaitán et al. 2018). It can survive at elevated temperatures and high salt concentrations, which otherwise act as physiological barriers to fungal infections (Casadevall et al. 2019; Jackson et al. 2019). As a haploid ascomycete, *C. auris* often displays exceptional resistance to major antifungals like azoles and common sterilization agents, rendering it a difficult pathogen to treat (Emara et al. 2015; Cadnum et al. 2017; Chowdhary et al. 2018). As an opportunistic pathogen, *C. auris* colonises skin and causes systemic infections, thereby posing threats to patients with other clinical conditions like diabetes mellitus, chronic renal disease, and more recently COVID-19 infections (de Cássia Orlandi Sardi et al. 2018; Rodriguez et al. 2020). *C. auris* emerged and evolved simultaneously as distinct geographical clades - South Asian (clade 1), East Asian (clade 2), South African (clade 3), South American (clade 4) and a potential fifth clade from Iran (Lockhart et al. 2017; Chow et al. 2019).The clades are separated by tens of thousands of single nucleotide polymorphisms (SNPs) but exhibit clonality within a clade (Lockhart et al. 2017). The mechanisms that underlie the sudden emergence and spread of *C. auris* as distinct geographical clades, though mostly unknown, represent rapid evolution modes in a fungal pathogen.

Both the pathogen and its host coevolve in nature to survive the evolutionary arms race. Chromosomal reshuffling serves to generate diversity in some predominantly asexual fungal pathogens (Sun et al. 2017; Guin, Chen, et al. 2020; Sankaranarayanan et al. 2020; Schotanus & Heitman 2020), thereby circumventing evolutionary dead ends. Chromosomal rearrangements and aneuploidy are also known to enhance drug resistance and virulence in primarily asexual fungi (Selmecki et al. 2008; Poláková et al. 2009; Legrand et al. 2019). Centromeres (*CEN*s) that appear as the primary constrictions on metaphase chromosomes, are emerging as a central hub of such chromosomal rearrangements contributing to karyotype diversity and speciation (Guin, Sreekumar, et al. 2020). Centromeres exhibit diversity in their properties like the length of centromeric chromatin, repeat/transposon content, and GC-richness. However, centromeric chromatin in most species is occupied by the *CEN*-specific histone variant CENP-A^Cse4^, that replaces canonical histone H3 in the centromeric nucleosomes and is regarded as the epigenetic hallmark defining *CEN* identity (McKinley & Cheeseman 2016; Yadav et al. 2018). Centromeric chromatin also provides foundation for assembling several multiprotein complexes to form the kinetochore. Dynamic interactions of spindle microtubules and kinetochores result in the precise segregation of sister chromatids in daughter cells during cell division. Centromere clustering near the nuclear periphery is a conserved feature across the fungal kingdom (Sanyal & Carbon 2002; Padmanabhan et al. 2008; Navarro-Mendoza et al. 2019; Fang et al. 2020). Due to spatial proximity, centromeres with homologous DNA sequences often participate in chromosomal rearrangements that result in chromosomal shuffling which can drive karyotype evolution and chromosome number alterations, contributing to the emergence of a new species (Guin, Chen, et al. 2020; Ola et al. 2020; Sankaranarayanan et al. 2020).

*C. auris* is a sister species of three multidrug-resistant pathogens, namely, *Candida haemulonii*, *Candida duobushaemulonii*, and *Candida pseudohaemulonii*. These species are also closely related to another human fungal pathogen, *Candida lusitaniae*, and together are classified under the *Clavispora/Candida* clade of the family Metschnikowiaceae (Order: Saccharomycetales) (Gabaldón et al. 2016; Muñoz et al. 2018). Centromeres are susceptible to breaks in other fungal pathogens and are likely to contribute to the vast karyotype diversity exhibited by *C. auris* (Chatterjee et al. 2016; Bravo Ruiz et al. 2019; Guin, Chen, et al. 2020; Sankaranarayanan et al. 2020). We believed that studying the centromere structure and function in the *C. haemulonii* species complex and associated species may reveal mechanisms/events underlying the rapid evolution of the multidrug resistant fungal pathogen *C. auris*. In this study, we identified centromeres in all four clades of *C. auris* and leveraged the information to locate centromeres in the *C. haemulonii* complex species. Functional identification of centromeres combined with comparative genome analysis in these group of species helped us propose that a centromere inactivation event from an ancestral species facilitated genome innovations that led to the clade-specific parallel evolution of *C. auris*.

## Results

### *C. auris* possesses small regional CENP-A^Cse4^-rich GC-poor, repeat-free centromeres

The histone H3 variant CENP-A^Cse4^ is exclusively associated with centromeric nucleosomes. The homolog of CENP-A^Cse4^ was identified in *C. auris*, using the *C. albicans* CENP-A^Cse4^ protein sequence as the query against the *C. auris* genome (GenBank assembly GCA_002759435.2). The putative *C. auris* CENP-A^Cse4^ protein is 136 amino acid long and shares a 72% sequence identity with the *C. albicans* homolog (C3_00860W_A) (supplementary fig. 1). Previous studies suggested that the haploid genome of *C. auris* is distributed in seven chromosomes (Muñoz et al. 2018). To locate centromeres on each chromosome, we constructed a strain CauI46 expressing Protein A-tagged CENP-A^Cse4^ from a clade 1 Indian isolate Cau46 (supplementary fig. 2A). Immunofluorescence staining using anti-Protein A antibodies revealed punctate localization of CENP-A^Cse4^ at the nuclear periphery, suggesting typical kinetochore clustering at interphase and mitotic stages of the cell cycle (fig. 1A). High amino-acid sequence similarities with other proteins of the CENP-A family and typical localization patterns of the clustered centromeres at the nuclear periphery confirmed that the identified protein is, indeed, CENP-A^Cse4^ in *C. auris*. To identify CENP-A^Cse4^ associated DNA sequences as centromeric chromatin on each chromosome of *C. auris*, we performed CENP-A chromatin immunoprecipitation (ChIP) followed by sequencing (ChIP-sequencing) in the strain CauI46. Sonicated genomic DNA-without antibodies was also subjected to high-throughput sequencing that served as the input DNA control. The CENP-A^Cse4^ ChIP-seq analysis identified a single-peak in each of the seven different scaffolds out of 15 scaffolds of the publicly available *C. auris* clade 1 reference genome assembly (fig. 1B). The CENP-A^Cse4^ enriched centromeric chromatin across chromosomes spans 2516 bp to 2908 bp, with an average length of 2727 bp (table 1). Further analysis of these regions suggests that CENP-A^Cse4^-enriched core centromere (*CEN*) loci in *C. auris* are largely devoid of ORFs and represent poly-(A) transcriptional cold spots (fig. 1C). To further confirm ChIP-seq results, ChIP-quantitative PCR (ChIP-qPCR) using specific primers was performed to measure CENP-A^Cse4^ abundance at *CEN*s compared to a non-centromeric genomic locus, ~200 kb away from *CEN4* (*far-CEN4*). The same centromeric and non-centromeric primer pairs (supplementary table 3) were used to assess the canonical histone H3 occupancy in the corresponding regions by histone H3 ChIP-qPCR analysis. As expected, histone H3 levels were significantly depleted at the *CEN*s compared to the *far-CEN* region (fig. 1D). Binding of CENP-A^Cse4^ to transcriptionally inert, histone H3-depleted loci of comparable length on different contigs strongly indicates that these genomic regions correspond to authentic centromeric chromatin.

**Table 1.**
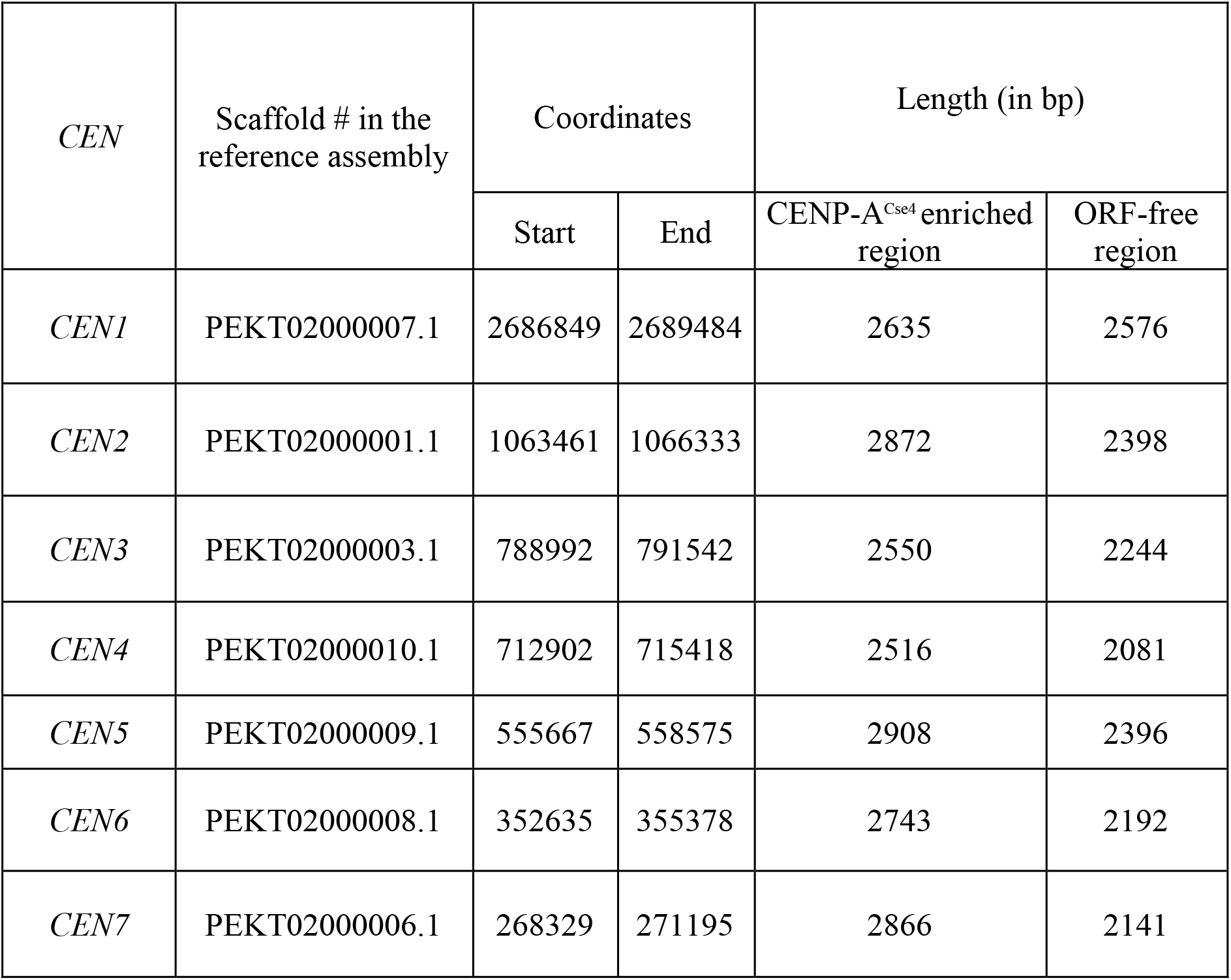
Centromere features in clade 1 isolate of *C. auris*

**Fig. 1.**
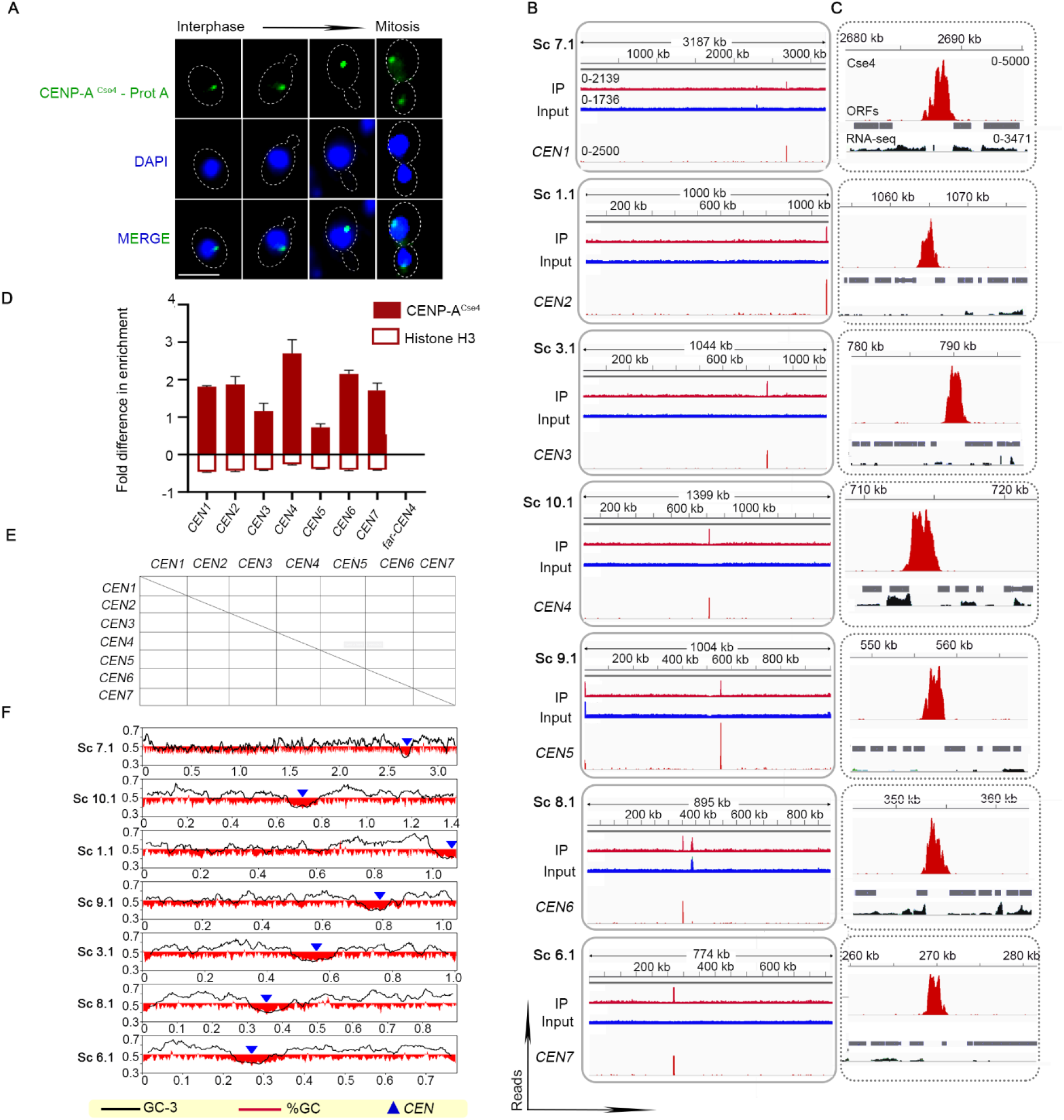
CENP-A^Cse4^-rich unique DNA sequences that are significantly depleted of histone H3 define small regional centromeres (*CEN*s) in each of seven chromosomes in *C. auris* clade 1. **A**, Indirect immunolocalization of Protein A-tagged CENP-A^Cse4^ (green) shows centromeres are clustered at the periphery of the nucleus stained with DAPI (blue) at various stages of the cell cycle. Scale bar, 3 μm **B**, CENP-A^Cse4^ ChIP-seq reads: Input-total DNA, IP-immunoprecipitated DNA, *CEN*-input subtracted from IP. **C**, Zoomed-in CENP-A^Cse4^ ChIP-seq peaks (red) along with ORFs (grey) and mapped RNA-seq reads (black). The peak values are indicated. **D**, Fold difference in CENP-A^Cse4^ and histone H3 enrichment at the *CEN*s compared to a control region (*far-CEN4*). qPCR values from three technical replicates are shown. The experiments were repeated thrice. Error bars indicate standard error of the mean (SEM). Statistical analysis: one-way ANOVA, **** P<0.0001 **E**, Dot-plot analysis reveals the absence of repeats and the unique nature of *CEN* DNA sequences **F**, *CEN* positions 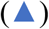 overlap with GC 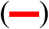 and GC-3 (–) scaffold minima. Coordinates (in Mb) are shown on the *x*-axis and %GC, on the *y*-axis.

Homology searches for *CEN* sequences among themselves and against the whole genome did not yield any significant results, suggesting that each DNA sequence underlying centromeric chromatin is unique and different. A dot-plot comparing each centromere DNA sequence against itself as well as other centromeric sequences suggested the unique nature of sequences and the absence of DNA sequence repeats in *C. auris* centromeres (fig. 1F). Searches for specific DNA sequence motifs also did not detect any, except the poly (A) and poly (T) stretches, which are present in all the seven regions, though not exclusive to the centromeres (supplementary fig. 2B). The presence of poly(A) stretches at all centromeres prompted us to analyse the GC-content of the *CEN* sequences identified. Two sequence features were investigated using the sliding window approach-GC content (the percentage of G and C residues in the scaffold in a sliding window of 5 kb, with a step size of 1 kb) and GC3 content (GC content at the third position of codons in the annotated ORFs, across the scaffolds). These studies revealed the overlap of *C. auris* centromere*s* with deep GC and GC3 troughs in all the scaffolds (fig. 1G).

At each of the seven centromeres in *C. auris*, core CENP-A^Cse4^ chromatin occupies the entire ORF-free region, often extending partially to the neighbouring centromere-proximal ORFs. By comparing the lengths of CENP-A^Cse4^-bound and the associated ORF-free regions in the previously characterized centromeres of Ascomycota, we observed that centromeric chromatin tends to possess a localized region within the gene-poor zones in species like *C. albicans* and *S. cerevisiae*. Exceptionally, the ratio of centromeric chromatin to the remaining ORF-free pericentric region in *C. auris*, similar to that of *C. lusitaniae*, is close to 1 (supplementary fig. 2C). Thus, *C. auris*, like *C. lusitaniae* seems to lack pericentric heterochromatin(Kapoor et al. 2015). We analysed RNA-seq data available for *C. auris* (SRR6900290, SRR6900291, SRR6900292, SRR6900293) to examine variations of gene expression at the centromere vicinity that might indicate the presence of pericentric heterochromatin. We could not detect any suppression of gene expression in the centromere neighbourhoods (supplementary fig. 2D, E), confirming that *C. auris*, like *C. lusitaniae*, possess pericentric heterochromatin-deficient (PHD) centromeres (supplementary fig. 2F). Pericentric heterochromatin formation is a concerted function of pericentric repeats, RNA interference machinery, chromodomain proteins, methyl transferases as well as histone deacetylases. However, these factors have a patchy distribution in the fungal kingdom (Bühler & Moazed 2007; Drinnenberg et al. 2009; Hickman et al. 2011; Alper et al. 2013). As expected, orthologs of many heterochromatin-forming proteins could not be detected in the *C. auris* and *C. haemulonii* complex species (supplementary table 4). Though orthologs of Dcr1 (the non-canonical Dicer protein) are present, Ago1 (Protein argonaute) and Rdp1 (RNA-dependent RNA polymerase) could not be detected in any of these ascomycetes.

### Clade-specific karyotype alterations in *C. auris* involve centromeres

Clinical isolates of *C. auris* have been primarily classified into four geographical clades, which exhibit differences in virulence, drug resistance, and genome plasticity. Having identified centromeres in a clade 1 isolate, we sought to identify centromere loci in other clades of *C. auris*. Are the centromeres and their neighbourhoods conserved in sequence and location across different geographical clades? To answer this, we predicted the putative centromere coordinates in clades 2, 3, and 4 of *C. auris* based on gene synteny, GC-content, and ORF-content using the available assemblies (GCA_003013715.2 of strain B11220 for clade 2, GCA_005234155.1 of strain LOM for clade 3, and GCA_008275145.1 of strain B11245 for clade 4) (fig. 2A). The predictions were experimentally tested using strains expressing CENP-A^Cse4^ - Protein A fusion proteins in each of these three clades. The predicted loci were enriched with CENP-A^Cse4^ and depleted of canonical histone H3 (supplementary fig. 3A, B, D, E, G, H). Like clade 1, all seven identified centromeres in each of the three clades overlap with GC- and GC3-troughs (supplementary fig. 3C, F, I)). Taken together, we identify small regional AT-rich centromere loci of all chromosomes in each of the four clades of *C. auris*.

**Fig. 2.**
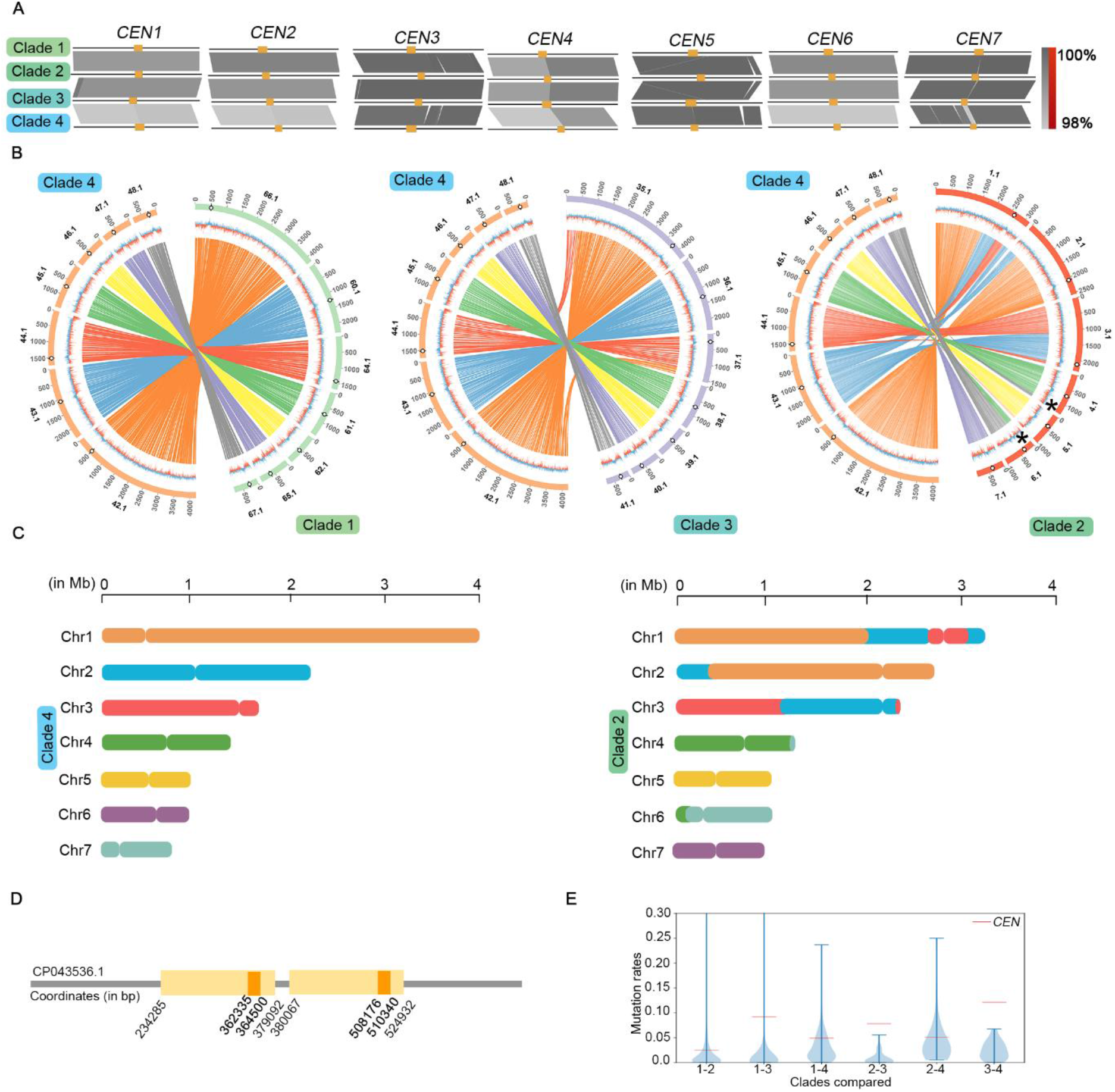
Chromosomal rearrangements resulted in an exclusive centromere relocation in clade 2. **A**, Diagram showing immediate *CEN* neighbourhood conservation (20 kb each to the left and right of *CEN*s, marked in orange) in each of the four clades. 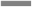 connects homologs; inversions, if present, are shown by 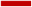. The sequence similarity is shown as a percentage. **B,** Circos plots showing synteny conservation between different clades. Scaffold numbers are shown on the outer-most track with empty circles marking centromere positions, GC-content is shown in the middle track (red, GC content below genome average and blue, AT content above genome average), and the inner-most track shows synteny blocks. A reciprocal translocation event in clade 2 is marked by *. **C**, Linear synteny plot showing *CEN* relocations in clade 2 with respect to those of clade 4. *CEN* positions in clades 1 and 3 are similar to clade 4 *CEN* locations. *CEN*s are shown as chromosomal constrictions. Chromosomes are drawn to scale, and chromosomal sizes are shown. **D,** Schematic depicting segmental duplication (yellow) in clade 2, placing two copies of the centromere sequences (orange) in the same chromosome in the assembly GCA_003013715.2. The scaffold number and the coordinates are shown. **E**, Violin plot depicting mutation rates at the centromere sequences, compared to intergenic regions in each pair of clades. Standard deviation for the mutation rates at intergenic region is shown, and the mutation rates at the centromeres are shown as Z-scores (difference from mean in units of standard deviation).

Next, we performed genome-wide comparisons using the publicly available chromosome-level assemblies of *C. auris* to study the involvement of centromeres in clade-specific rearrangements, if any. From MLST analysis based on *RPB2* (Prakash et al. 2016), *TUB2*, and *EFB2* gene sequences, we observed that strain A1, isolated in China (SRS4986047), belongs to clade 3 and strain CA-AM1 (SRS7388889), isolated in Italy, belongs to clade 1. Centromere locations in these isolates were also identified. Centromere coordinates of all the isolates analysed are listed in supplementary table 5. Based on the presence of centromeres and syntenic regions shared with CA-AM1, we propose the merger of scaffold PEKT02000002.1 to PEKT02000001.1, PEKT02000005.1 to PEKT02000003.1, and PEKT02000004.1 to PEKT02000007.1 in the current reference assembly of clade 1 to fill the gaps and construct an improved assembly.

Both GCA_014673535.1 (for strain CA-AM1) and GCA_014217455.1 (for strain A1), being complete assemblies with seven contigs, were used as clade 1 and clade 3 assembly, respectively, for genome-wide comparisons. All combinations of pair-wise comparisons revealed inter-clade chromosomal changes in *C. auris*. Representative images using clade 4 (GCA_008275145.1) assembly as the reference is shown in fig. 2B. Centromeres were numbered from 1 to 7 in the clade 4 assembly based on the decreasing sizes of the chromosomes harbouring them. Centromeres of clades 1, 2, and 3 were numbered based on synteny with clade 4 *CEN*s. Cross-clade comparisons revealed the genome of clade 2 to be the most rearranged one compared to the other three clades, as reported previously (Muñoz et al. 2019) (fig. 2B). Five out of seven chromosomes in clade 2 had undergone chromosomal rearrangements, and two of these rearrangements in chromosomes 1 and 3 involve synteny breaks adjacent to the centromeres. These structural changes resulted in centromere relocations in clade 2 compared to other clades, generating significant karyotype alterations (fig. 2C). We also detected a segmental duplication in the clade 2 reference assembly (GCA_003013715.2). Duplication of a 145 kb fragment in contig000006 in the clade 2 assembly places two copies of the centromere region on the same contig, separated by 144 kb (fig. 2D).

Centromeres were earlier shown to be the most rapidly evolving loci in two closely related species of the CTG-Ser1 clade: *Candida albicans* and *Candida dubliniensis* (Padmanabhan et al. 2008). A similar genome-wide analysis among the clades of *C. auris* suggested that centromeres exhibit high incidence of substitution mutations compared to the intergenic regions of the genome. This is true for all the clades, though the rates of sequence changes are different (fig. 2E, supplementary table 6). Hence, a geographical clade-specific accelerated evolution of centromere sequences in the same species is evident from these analyses.

### *C. haemulonii* and related species share centromere properties with *C. auris*

The size of the *C. auris* genome is 12.2-12.4 Mb that falls in the same range with genomes of phylogenetically related, multidrug-resistant, pathogenic species *C. haemulonii*, *C. duobushaemulonii*, and *C. pseudohaemulonii* of sizes 13.3 Mb, 12.6 Mb, and 12.6 Mb, respectively (based on corresponding NCBI GenBank assemblies-see Methods). Since all these species of the *C. haemulonii* complex share similar biochemical properties, the misidentification of species in clinics is quite common. Gene synteny around the *CEN* neighbourhoods in these species is conserved compared to *C. auris*, enabling the prediction of *CEN* coordinates (fig. 3A, supplementary fig. 4A, E). The predicted *CEN* regions were also found to be histone H3-depleted and overlapping with scaffold GC-and GC3 minima (fig. 3B, 3C, supplementary fig. 4B-D, F-H), suggesting that these are the bona fide *CEN*s. The identified regions are largely free of ORFs and have lengths comparable to those of *C. auris CEN*s (supplementary table 7). Comparisons utilizing the available chromosome level assembly of *C. duobushaemulonii* revealed that this species is closer to clades 1, 3, and 4 than clade 2 of *C. auris* (supplementary fig. 5A-C), further corroborating the distinctiveness of clade 2, isolates of which are usually drug sensitive.

**Fig. 3.**
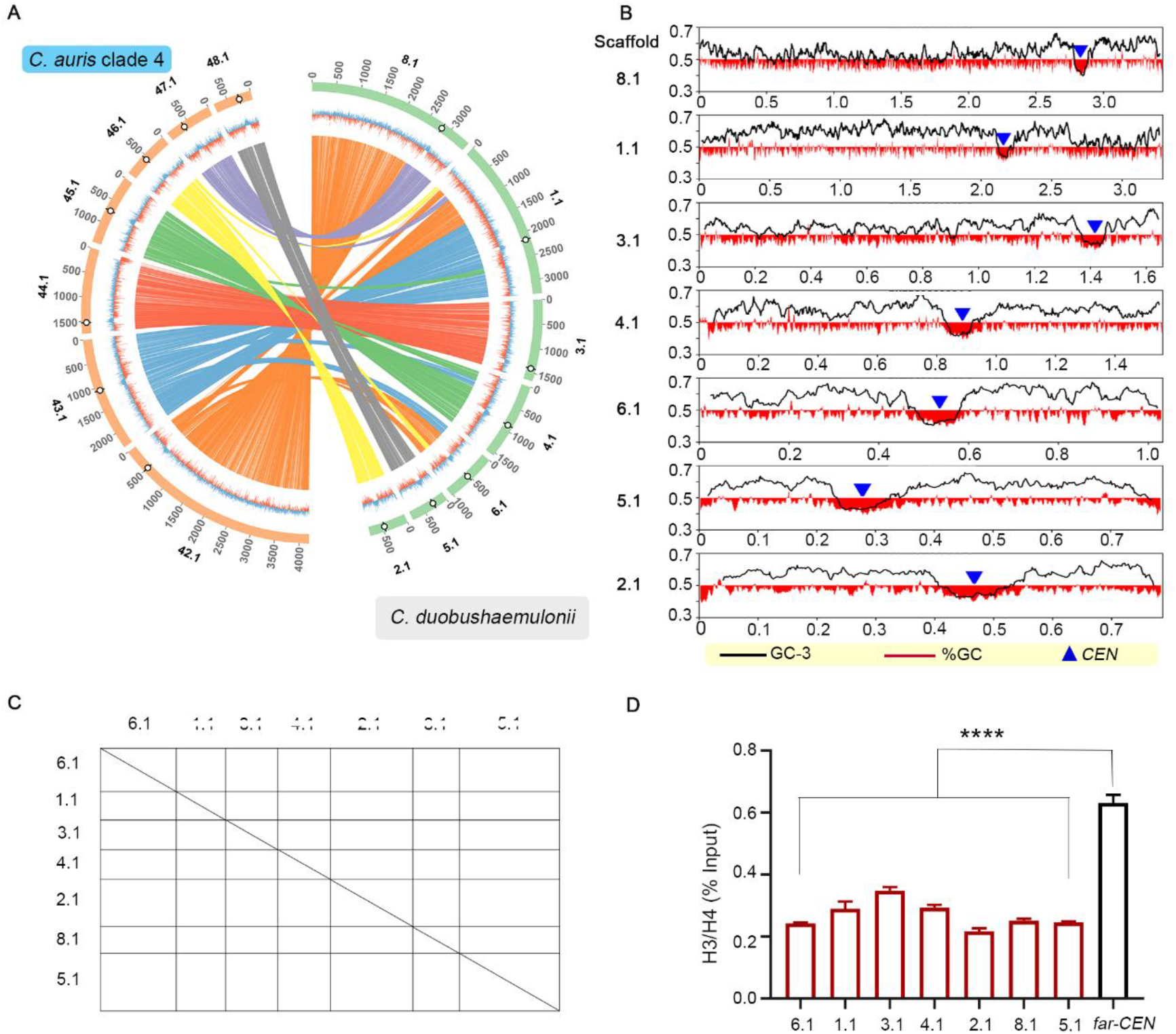
Conservation of centromere properties of the *C. haemulonii* complex species including C. *auris*. **A**, Loci in *C. duobushaemulonii* syntenic to *C. auris CENs*. The outer-most track of the circos plot depicts genome scaffolds with empty circles marking *CEN* locations, the middle track depicts %GC (red-GC content below genome average, blue-AT content above genome average), and the inner-most track shows synteny blocks. **B**, *CEN* positions 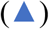 overlap with GC-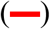 and GC-3 (–) minima. Coordinates (in Mb) are shown on the x-axis and %GC, on the y-axis. **C**, Dot-plot establishing the repeat-free and unique nature of centromere sequences in *C. duobushaemulonii*. The scaffold numbers are shown. **D**, Depletion of histone H3 at *CEN*s on different scaffolds (shown on the x-axis), compared to a non-centromeric control region (far-*CEN*). qPCR values from three technical replicates, represented as percent input, are shown. The experiments were performed thrice, with similar results. Error bars indicate standard error of the mean (SEM). Statistical analysis was done using one-way ANOVA (**** P<0.0001).

### A centromere inactivation event accounts for the chromosome number alteration between *C. lusitaniae* and *C. auris*

*Candida lusitaniae*, another opportunistic pathogen, is classified under the *Clavipora/Candida* clade of Metschnikowiaceae and is phylogenetically close to *C. auris* (fig. 4A). It was previously reported to have eight AT-rich short regional *CEN*s made up of unique DNA sequences (Kapoor et al. 2015). On the other hand, we report that *C. auris* has seven functional *CEN*s identified in this study. To trace the events that led to the chromosome number reduction during the divergence of these two species, we compared the gene synteny across the centromeres in *C. lusitaniae* and *C. auris*. Though the genomes are highly rearranged (supplementary fig. 5D), we found that the gene synteny around centromeres is conserved between the two species. Intriguingly, chromosome 8 of *C. lusitaniae* was rearranged as three distinct fragments that fused with other chromosomes of *C. auris*. As a result, two *C. lusitaniae* centromeres (*ClCEN2* and *ClCEN8*) were mapped to the same *C. auris* chromosome, based on synteny analysis (fig. 4B). ChIP-seq analysis revealed *CEN2* to be functional in *C. auris* out of the two regions as CENP-A^Cse4^ is recruited only at *CEN2*. This observation illustrates a clear example of “evolution in progress” as the region corresponding to *C. lusitaniae CEN8* becomes non-functional in *C. auris* despite gene synteny conservation between the two species around this region. *ClCEN8*, the functional centromere of chromosome 8 in *C. lusitaniae*, spans a region of around 4.5 kb, while the average centromere length is 4.3 kb. The size of the corresponding syntenic regions of the inactivated centromere (in*CEN*) is 1.1 kb in *C. auris*. In comparison, the functional centromeres of the same species have an average length of 2.7 kb. We posit that the significant, centromere-specific attrition of DNA sequence accompanied by the reduction of AT-content resulted in the centromere inactivation in *C. auris* (fig. 4C). Analysis at the sequence level reveals mutation rates at the in*CEN* to be intermediate of that of centromeres and intergenic regions, further suggesting a “transition from centromeric to intergenic region” (supplementary table 6).

**Fig. 4.**
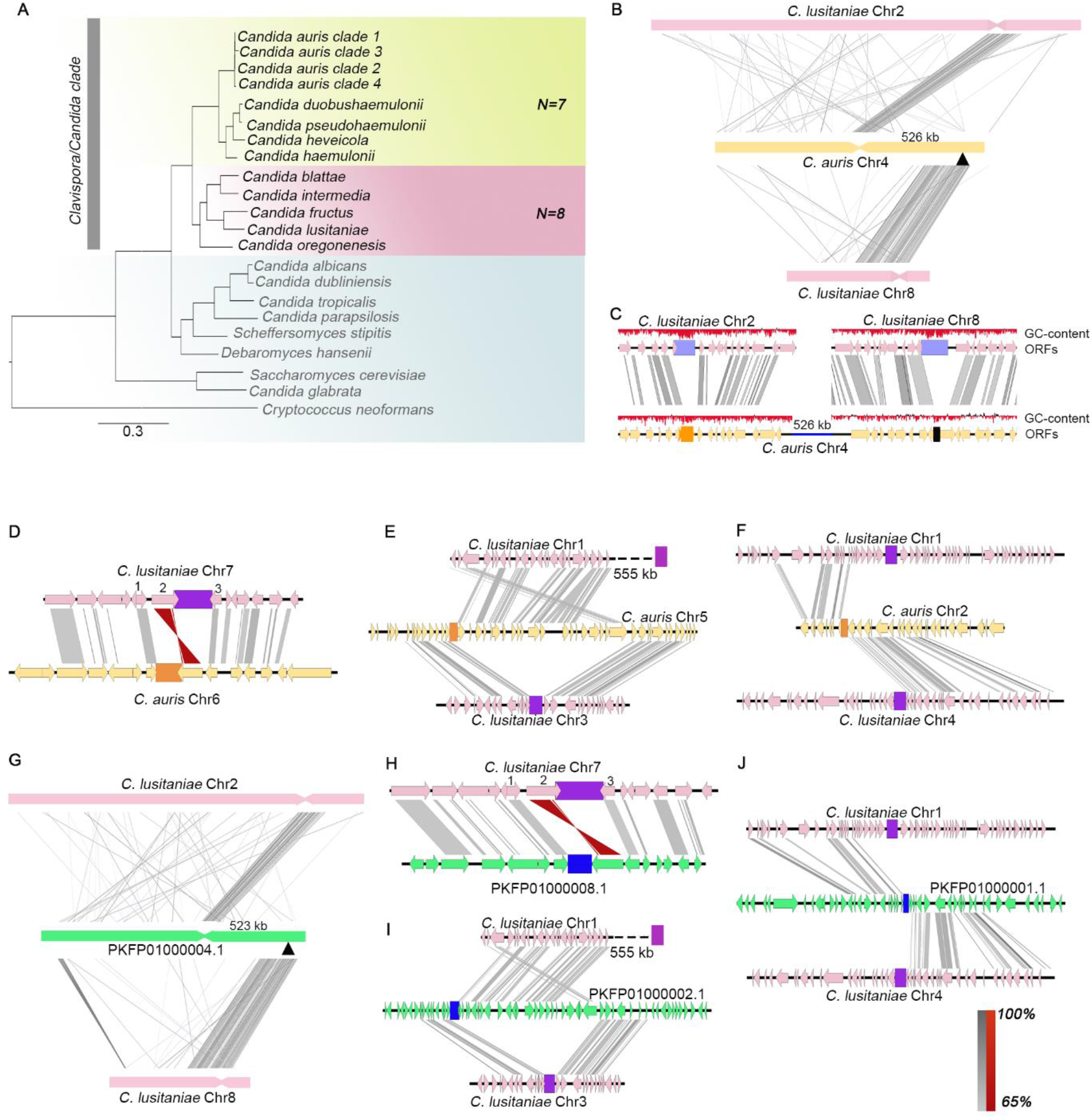
CEN-inactivation mediated chromosome number variation in *C. auris* and *C. duobushaemulonii*. **A**, Phylogenetic tree depicting the relatedness of *C. auris* geographical clades and other member species of the *Clavispora/Candida* clade. Other species in Ascomycota with characterised/predicted centromeres are shown. *Cryptococcus neoformans* (Basidiomycota) is shown as the outgroup. The two chromosome number states detected in *Clavispora/Candida* clade are represented by *N*=7 and *N*=8. **B**, Chromosome-level view depicting the mapping of *C. lusitaniae CEN2* and *CEN8* onto a single scaffold in *C. auris*. 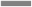 connects homologs; inversions, if present, are shown by 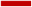. Inactive *CEN* (in*CEN*) is shown as ▲. The sequence similarity of homologs is shown in the key as a percentage. **C**, ORF-level view showing sequence loss and subsequent loss of AT-content at in*CEN*. **D**, Pericentric inversion in *C. auris* changing the positions of ORFs 1,2, and 3, with respect to the *C. lusitaniae* centromere 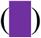. *C. auris CENs* are shown as 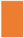. **E**, Rearrangement involving *CEN*-proximal synteny breaks separating the two synteny blocks on the same chromosome. **F**, Synteny breakpoint mapped to the centromere location in *C. auris* chromosome 2. **G**, *CEN* inactivation, **H**, pericentric inversion **I, J**, synteny breaks and rearrangements in *C. duobushaemulonii. CENs* are marked by 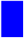. Inactive *CEN* is shown as ▲.

A distinct *CEN*-associated structural change observed in *C. auris*, compared to the syntenic *CEN* in *C. lusitaniae*, is a pericentric inversion altering the relative positions of three ORFs (fig. 4D). In addition to the presence of in*CEN*, five centromere regions in *C. lusitaniae* have syntenic centromeres in *C. auris*. The remaining two, identified through CENP-A^Cse4^ ChIP-seq, are located at synteny breakpoints and hence, are involved in chromosomal rearrangements. The immediate ORFs flanking *CEN3* in *C. lusitaniae* are conserved in *C. auris* but are separated by a length of 55 kb. The centromere is located adjacent to one of the synteny blocks, resulting in partial synteny conservation (fig. 4E). We also mapped a synteny breakpoint at the centromere on chromosome 2 of *C. auris*. The ORFs on either side of the

*C. auris CEN2* maps to different chromosomes in *C. lusitaniae* (fig. 4F). The same patterns were observed in *C. haemulonii*, *C. duobushaemulonii*, and *C. pseudohaemulonii*, where sequences syntenic to *ClCEN8*-flanking blocks map to the same scaffold bearing *ClCEN2* synteny regions (fig. 4G, supplementary fig. 6A, B). The region corresponding to *ClCEN8* has undergone differential sequence attrition in these species, resulting in reduced sequence length (840 bp in *C. haemulonii*, 361 bp in *C. duobushaemulonii*, and 496 bp in *C. pseudohaemulonii*) as observed in *C. auris* in*CEN*. *CEN*-specific sequence loss has also resulted in the reduction of AT-content in these species. *CEN*-associated inversions and synteny breakpoints in these species are also identical to those in *C. auris* (fig. 4H-J, supplementary fig. 6C-H). The typical patterns of *CEN*-associated changes in *C. auris* and other species of the *C. haemulonii* complex suggest that these events must have occurred in an immediate common ancestor before species divergence.

### Putative small regional, AT-rich centromeres identified in other species of the *Clavispora/Candida* clade

Around 40 ascomycetous species are classified under the *Clavispora/Candida* clade of Metschnikowiaceae (Daniel et al. 2014). To explore the centromere properties in the *Clavispora/Candida* clade, we attempted *CEN* identification in other species for which genome assemblies are available (fig. 4A). We could locate putative centromeres in several fungal species of the *Clavispora/Candida* clade of Metschnikowiaceae based on the conserved gene synteny and other conserved centromere properties of *C. auris* and *C. lusitaniae* as references (supplementary table 8-12). Two possible chromosome number states were detected in the *Clavispora/Candida* clade, and the analysed genomes were classified into two groups – a) species which have eight AT-rich putative centromeric loci of comparable sizes, and b) species with seven AT-rich putative centromeric loci with an eighth locus that had undergone sequence loss despite synteny conservation around the orthologous but presumably inactivated centromere locus. *C. lusitaniae* has eight AT-rich, ORF-free centromeres of comparable lengths. *Candida fructus* was found to possess eight loci syntenic to each of the eight centromeres in *C. lusitaniae*. The identified regions are also depleted of ORFs, are GC-poor, and harbour GC-skews like *C. lusitaniae* centromeres (supplementary fig. 7). Each of *C. auris*, other species of the *C. haemulonii* complex, and *Candida heveicola* has seven ORF-free loci, which are GC-poor. The eighth locus, though syntenic to *CEN8* of *C. lusitaniae*, has undergone sequence attrition in each of them and is likely to be inactive, like the *inCEN* of *C. auris*. We could identify loci in other related species, including *Candida intermedia, Candida blattae*, and *Candida oregonensis* syntenic to each of the seven centromeres of *C. auris*. All the predicted regions are ORF-free, AT-rich, and constituted by unique, repeat-free sequences (supplementary fig. 8A, B).We also identified an eighth locus syntenic to *C. lusitaniae CEN8* in these species. Unlike the in*CEN* in *C. auris* with a drastically reduced sequence length, the eighth locus is of similar size as other predicted centromeres in these three species (supplementary fig. 8A, C). The conservation of sequence length suggests that they may have eight functional centromeres. Exceptionally due to a possible assembly error, two putative centromeres identified in *C. intermedia* map to the same scaffold. Our *in silico* analyses collectively suggest the existence of two chromosome number states and remarkably similar centromere properties shared by these closely related organisms of the *Clavispora/Candida* clade. While all these putative *CEN* loci show similar gene synteny, ORF-abundance, sequence length, and GC-content, further experimental validation is required before assigning them as authentic *CEN* loci of the respective organisms.

### Clade 2 of *C. auris* follows a unique evolutionary trajectory

We posit that *C. lusitaniae* and *C. fructus* might have shared an immediate common ancestor CA1 with eight functional *CEN*s, one on each chromosome (*N*=8). Chromosomal rearrangements placed regions syntenic to *CEN2* and *CEN8* of these two species on the same chromosome in the *C. haemulonii* complex species as well as three clades (clades 1, 3, and 4) of *C. auris*, out of which *ClCEN2* is active, and *ClCEN8* is inactive (in*CEN*) (fig. 5A). This finding indicates the existence of an immediate common ancestor (*N=7*), CA2, with a *ClCEN2-inCEN* configuration shared by *C. auris* and other species of the *C. haemulonii* complex. Synteny analyses enabled us to reconstruct (fig. 5A) *CEN*-based ancestral genomes of the immediate common ancestors of *C. lusitaniae-C. fructus* and *C. haemulonii* complex-*C. auris*, representing chromosome number states of *N=8* and *N=7*, respectively. We also hypothesize parallel evolution of the geographical clades of *C. auris*, at different time scales, diverging from a common ancestor CA3, which was derived from the ancestor CA2. Out of the four clades, clade 2 has a remarkably rearranged genome. The location of in*CEN* serves as a useful index for representing interclade differences.The synteny block containing *C. lusitaniae CEN8* is conserved in *C. haemulonii*, *C. pseudohaemulonii*, *C. duobushaemulonii* as well as in *C. auris* clades 1,3, and 4. The genes in the block are found distributed in two chromosomes in clade 2, indicating that a break occurred within the block, followed by a downstream reciprocal translocation event (supplementary table 13, fig. 2B). The terminal chromosomal translocation (TCT) event in which Chr4 and Chr7 of CA3 exchanged chromosome ends might have repositioned in*CEN* resulting in a *ClCEN5-inCEN* configuration (fig. 2B, fig. 5B), exclusive to clade 2. This structural change further confirms the divergence of clade 2 from the common ancestor CA3 along a different evolutionary trajectory (fig. 5C). On analysing the whole genome synteny conservation, we observed that clades 1,3, and 4 are closer to *C. duobushaemulonii* (supplementary fig. 5), supporting the inference that clade 2 is unique. Also, the conservation of the *C. lusitaniae CEN8*-containing synteny block among the *C. haemulonii* complex species and all the *C. auris* clades except clade 2 suggests that each of these species is phylogenetically closer to *C. lusitaniae* than clade 2. These observations are in disagreement with an alternative model of clade 2 being the ancestral unique strain where the event leading to chromosome number reduction happened in which case clade 2 would have shared higher similarity with *C. lusitaniae*. Other rearrangements causing *CEN* relocations provide additional lines of evidence for the clade-specific divergence.

The inferred genomes can serve as references to trace *CEN*-associated rearrangements in other related species of the *Clavispora/Candida* clade. Other *CEN*-associated structural changes observed between *C. lusitaniae* and *C. auris* have an uneven distribution across the member species, indicating that *C. intermedia*, *C. blattae*, and *C. oregonensis* are likely to be transitional species connecting the two immediate common ancestors (supplementary table 14).

**Fig. 5.**
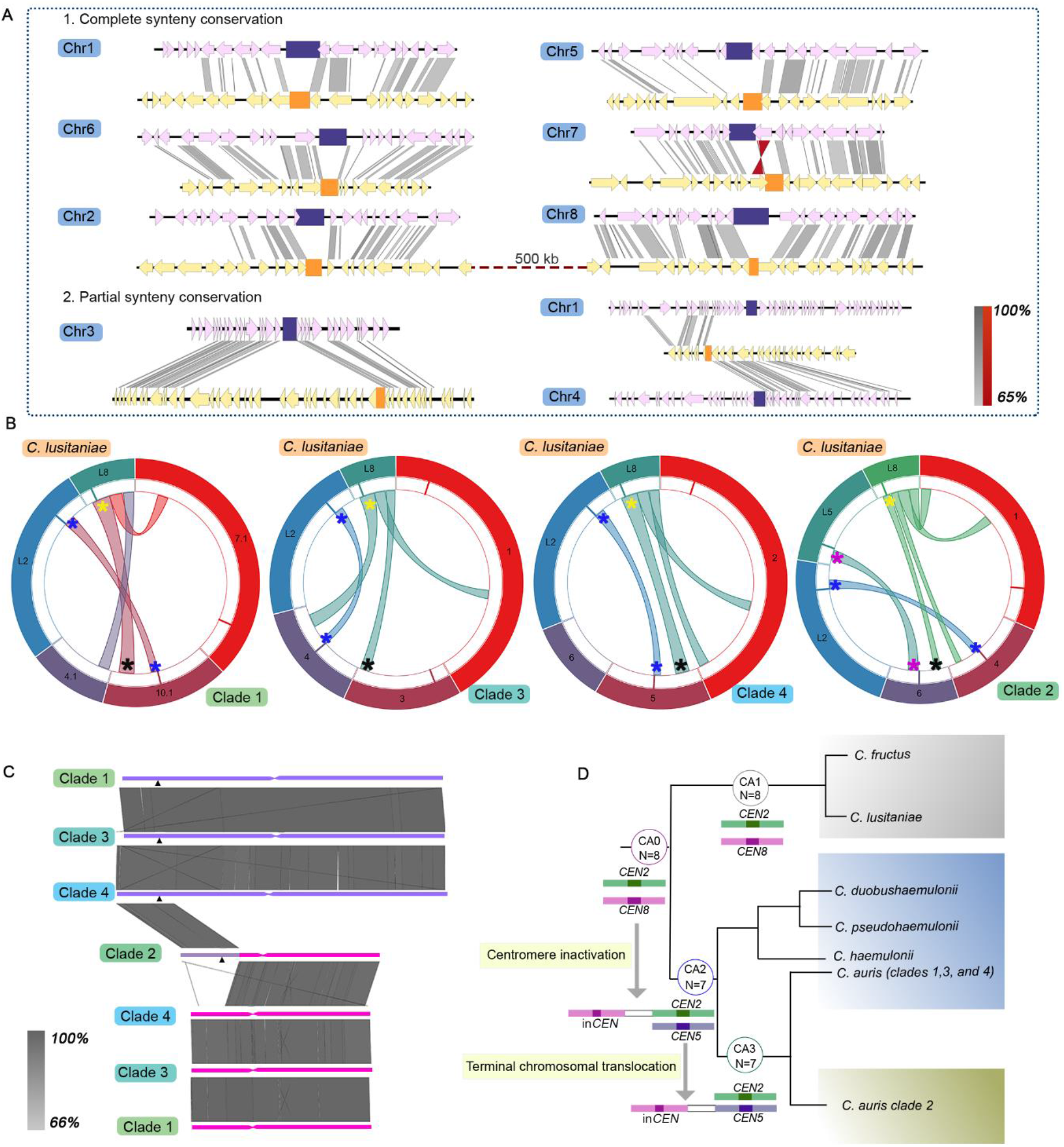
*C. auris* clade 2 evolved via a unique evolutionary trajectory. **A**, Representative genome of the common ancestor of *C. auris* and the *C. haemulonii* complex species (*N*=7), depicting chromosomal rearrangement patterns with respect to the common ancestor of *C. lusitaniae and C. fructus* (*N*=8). *CEN*s in the common ancestor (N=8) are shown as 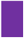, and *CEN*s in the common ancestor (*N*=7) are shown as 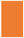. Homologs are connected by 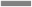 and inversions, if present, are shown as 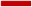. The sequence similarity is shown as a percentage in the key. **B**, *C. lusitaniae CEN2* (on Chr2, L2) and *CEN8* (on Chr8, L8) map to the same scaffold in *C. auris* clades 1,3 and 4, whereas *CEN5* (on Chr5, L5) and *CEN8* map to the same scaffold in *C. auris* clade 2. Corresponding genomic scaffolds are shown in the outer-most track, and the synteny blocks are depicted in the inner-most track. The inactive centromere is marked in black and the corresponding active centromere in yellow. **C**, Terminal chromosomal translocation event resulting in the relocation of in*CEN* (▲) in clade 2. Constrictions mark *CEN*s syntenic to *C. lusitaniae CEN2* and *CEN5*. Sequence similarity is shown as a percentage in the key. **D,** A *CEN*-based model tracing the event of centromere inactivation in the common ancestor CA0, resulting in chromosome number reduction in CA2, while CA1 maintains the chromosome number of 8. CA2 represents the common ancestor of *C. auris* and *C. haemulonii* complex, and CA3 is the common ancestor of all *C. auris* clades. A TCT event further repositions the inactive *CEN* in clade 2, representing the unique evolutionary trajectory of *C. auris* clade 2. *CEN*s are numbered using *C. lusitaniae* as the reference.

## Discussion

Centromere identification revealed a typical centromere landscape in multiple species of the *Clavispora/Candida* clade - small regional *CEN*s constituted by AT-rich unique sequences and embedded in ORF-free regions that are devoid of any detectable pericentric heterochromatin, DNA motifs or repeats. These closely related species either contain seven chromosomes or eight chromosomes. We propose that a centromere inactivation event in a common ancestor with eight chromosomes led to this diversity. The inactive centromere, in a pseudo-dicentric chromosome that might have formed at an intermediate stage, underwent substantial but differential attrition of centromere DNA sequence. This process might have played a crucial role in the emergence of multiple species with seven chromosomes. Inactivation of centromere function mediated by DNA sequence has been suggested previously (Jäger & Philippsen 1989; Gordon et al. 2011; Lhuillier-Akakpo et al. 2015). Several synteny breakpoints mapped to the identified centromeres, when compared with representative species of the eight-chromosome state, add to the growing evidence that suggests centromeres as a hub of fragility (Simi et al. 1998; Kim et al. 2013) and downstream chromosomal rearrangements. Spatial proximity of clustered centromeres in fungal species with homogenized centromere DNA sequences facilitates intercentromeric recombination, possibly mediated by replication fork stalling and higher chances of double-stranded breaks, thus contributing towards karyotype evolution (Greenfeder & Newlon 1992; Aze et al. 2016; Guin, Chen, et al. 2020). The role of AT-rich sequences and poly (A) stretches in these events, owing to their melting features and potential propensity to form non-B DNA, warrants further study as centromeres in many fungal species coincide with GC-or GC-3 troughs (Zhang & Freudenreich 2007; Lynch et al. 2010; Navarro-Mendoza et al. 2019; Yadav et al. 2019; Sankaranarayanan et al. 2020; Talbert & Henikoff 2020).

Whole chromosome and segmental aneuploidy are correlated with drug resistance in other fungal pathogens (Kwon-Chung & Chang 2012). The *C. auris* genome is known to be highly plastic (Bravo Ruiz et al. 2019). Mapping of centromere loci should help trace genomic rearrangement events that possibly contribute to drug resistance or virulence in different clinical isolates. Centromere sequences in different geographical clades were found to evolve rapidly and differentially than the rest of the genome, suggesting that centromeres are potential candidate loci to study evolutionary trajectories emerging within a species. *C. auris* clade 2 has the most rearranged genome and consists of atypical isolates that differ from the other clades in terms of drug tolerance as well as pathogenicity (Iguchi et al. 2019; Muñoz et al. 2019; Sekizuka et al. 2019). The unique nature of centromere sequences can be used for accurate species-level and clade-level identification.

In this study, we reveal that the genome of clade 2 differs from the rest of the clades in the position of orthologous centromeres on the chromosomes and the location of the inactive centromere. Chromosome-level comparisons also reveal that clade 2 is more diverged from *C. duobushaemulonii* than the other clades. These observations directed us to conclude that *C. auris* clades diverged from a common ancestor that shares ancestry with the *C. haemulonii* complex species, and from which clade 2 diverged along a different trajectory during the parallel evolution of the geographical clades. Drastic karyotype alterations, evident from the centromere and inactive centromere locations are likely to have contributed to the distinctiveness of *C. auris* clade2, compared to other clades and the *C. haemulonii* complex species. Ascomycetous pathogens like *C. albicans* and *C. glabrata* exist as clades that exhibit geographical specificity and clade-specific phenotypic features (Dodgson et al. 2003; Soll & Pujol 2003). Rare or no interclade recombination is observed in these species, and little is known about the genomic rearrangements or the variations at centromeres operating at the clade-level, which can, in turn, affect the recombination frequency.

We conjecture that such centromere-associated clade-specific differences might not be restricted to *C. auris*. Further exploration of centromere sequences and associated structural changes within a species and species complexes will yield deeper insight into the role of centromeres in generating diversity in primarily asexual fungi.

## Materials and Methods

### Strains, media, and growth conditions

Strains of various *Candida* species used in the study (listed in supplementary table 1) were grown in YPD (1% Yeast Extract, 2% Peptone, and 2% Dextrose) at 30 °C. The identity of the strains was confirmed by amplification and sequencing of the internal transcribed spacer (ITS) and D1/D2 regions, followed by BLAST analysis (http://www.ncbi.nlm.nih.gov/BLAST/Blast.cgi). The clade-status of different *C. auris* isolates used was confirmed by amplifying and sequencing regions of three housekeeping genes (*TUB2*, *EFB1*, and *RPB1*) harbouring polymorphic sites (supplementary table 2).

### Construction of *C. auris* strain expressing CENP-A^Cse4^-Protein A fusion protein

The homolog of CENP-A^Cse4^ in *C. auris* was identified by BLAST using *C. albicans* CENP-A^Cse4^ sequence as the query against the *C. auris* genome. It was distinguished from the canonical histone H3 sequences by confirming the presence of CENP-A^Cse4^ - specific amino acid residues (Keith et al. 1999). For tagging CENP-A^Cse4^ with Protein A at the C-terminus, approximately 900 bp and 800 bp were used as upstream and downstream sequences, respectively to construct the tagging cassette. The 900 bp fragment (including the complete ORF and native promoter sequence) was amplified from the genomic DNA and cloned as a *Kpn*I-*Sac*I fragment in the pBS-TAP-NAT plasmid. The downstream sequence was cloned as a *Spe*I-*Not*I fragment. The 3.7 kb tagging construct, as a *Kpn*I-*Not*I fragment, was used to transform Cau46R. The transformation of the strains was performed as previously described(Sanyal et al. 2004). Nourseothricin (Jena Bioscience) was added at a concentration of 100 μg/ml in the media for selecting transformants. The colonies obtained were subcultured in presence of nourseothricin and integration of the tagging construct in NAT^+^ transformants was confirmed by PCR.

### Western Blotting

Cells were grown overnight in YPD till mid-log phase, and 3 OD equivalent cells were harvested for protein lysate preparation. The cells were suspended in 400 μL of ice-cold trichloroacetic acid (12.5%), vortexed briefly, and stored at −20°C overnight. The samples were later thawed and pelleted by centrifugation at 14,000 rpm at 4 °C for 10 min. The pellets were washed twice with 400 μL of ice-cold acetone (80%), air-dried, suspended in an appropriate volume of lysis buffer (0.1 M NaOH and 1% SDS), and boiled for 10 min. The proteins in the lysate were separated on 12% polyacrylamide gels. The separated samples were transferred onto nitrocellulose membranes, which were then probed with anti-Protein A antibodies (Sigma, Cat No: P3775, 1:5000 dilution in 2.5% skim milk powder (w/v in 1x PBS)) and HRP-conjugated goat anti-rabbit secondary antibodies (Abcam, 1:10000 dilution in 2.5% skim milk powder (w/v in 1x PBS)). The blots were developed using Chemiluminescence Ultra Substrate (Biorad) and imaged using the VersaDoc system (Biorad).

### Preparation of spheroplasts

Cells were grown in 50 ml YPD till OD_600_ = 0.8 and washed with water by centrifugation at 3000 rpm for 5 min. Cells were then incubated in 10 mL of 2-mercaptoethanol solution (5% in water, Himedia, Cat No. MB041) for 1 h at 30°C at 180 rpm. The cells were pelleted, washed, and resuspended in SCE buffer (1M sorbitol, 100 mM sodium citrate, 10 mM EDTA at pH 8.0). Lysing enzyme from *Trichoderma harzianum* (Sigma, Cat No. L1412) was added at a concentration of 2.5 mg/ml, and the suspension was incubated at 37°C at 80 rpm for 2 h. The cells were examined under a microscope to determine the proportion of spheroplasts in the suspension. The prepared spheroplasts were further processed based on the corresponding experimental design.

### Indirect Immunofluorescence

*C. auris* CENP-A^Cse4^-Protein-A strain was inoculated to 1% (v/v) from an overnight culture and was grown till OD_600_ = 0.8. The cells were fixed by adding formaldehyde to a final concentration of 1% for 15 min. Spheroplasts were prepared from the fixed cells (as described above), washed with 1x PBS, and diluted in 1x PBS to a density appropriate for microscopy. Slides for microscopy were washed and coated with poly L-lysine (10 mg/mL). Twenty microlitres of the diluted cell suspension were added onto slides and incubated at room temperature for 5 min. The suspension was aspirated, and the slide was washed to remove unbound spheroplasts. The slide was treated with ice-cold methanol for 6 min, followed by ice-cold acetone for 30 sec. Blocking solution (2% non-fat skim milk powder in 1x PBS) was added to each well, and the slide was incubated for 30 min at room temperature. The blocking solution was aspirated, and rabbit anti-Protein A antibodies (Sigma, Cat No. P3775, dilution 1:1000) were added. The slide was incubated in a wet chamber for 1 h. The antibodies were aspirated, and the slide was washed 15 times, incubating the slide for 2 min for each wash. Secondary antibodies were added (Alexa flour 568 goat anti-rabbit IgG, Invitrogen, Cat No. A11011, dilution 1:1000). The slide was incubated in the dark in a wet chamber for 1 h at room temperature. The washes were repeated, and the mounting medium (70% glycerol with 100 ng/ml DAPI) was added. Clean coverslips were mounted onto the wells, and the slides were imaged using an inverted fluorescence microscope (Zeiss Axio observer, Plan Apochromat, 100X oil). Images were processed using Zeiss ZEN system software and ImageJ.

### Chromatin immunoprecipitation (ChIP)

*C. auris* CENP-A^Cse4^-Protein-A strain was inoculated to 1% (v/v) from an overnight culture, grown till OD_600_ = 1.0 and crosslinked by the addition of formaldehyde to a final concentration of 1% for 15 min. Quenching with 0.135 mM Glycine for 5 min was followed by preparation of spheroplasts (as described above). Following buffers were used to wash the prepared spheroplasts: 1x PBS (ice-cold), Buffer-1 (0.25% Triton X-100, 10 mM EDTA, 0.5 mM EGTA, 10 mM Na-HEPES at pH 6.5), and Buffer-2 (200 mM NaCl, 1 mM EDTA, 0.5 mM EGTA, 10 mM Na-HEPES at pH 6.5). One mL lysis buffer (50 mM HEPES at pH 7.4, 1% Triton X-100, 140 mM NaCl, 0.1% Na-deoxycholate, 1 mM EDTA) was added to the pellet obtained after the final wash, along with Protease inhibitor cocktail (1x). The resuspended spheroplasts were sonicated to obtain chromatin fragments in the size range of 100-400 bp. The lysate was cleared by centrifugation at 14,000 rpm for 10 min at 4 °C. One-tenth of the lysate volume was separated to be used as the input DNA. The remaining lysate was divided into two equal fractions: Anti-Protein-A antibodies were added to one of the fractions (IP fraction) at a 20 μg/mL concentration. The other fraction served as the antibody-minus control. Both the fractions were incubated overnight on a rotaspin at 4 °C. Protein-A Sepharose beads were added, and the samples were incubated on a rotaspin at 4 °C for 6 h. This was followed by collecting the beads by centrifugation and sequential washes with the following buffers: twice with 1 mL low salt wash buffer (0.1% SDS, 1% Triton X-100, 2 mM EDTA, 20 mM Tris at pH 8.0, 150 mM NaCl), twice with 1 mL high salt wash buffer (0.1% SDS, 1% Triton X-100, 2 mM EDTA, 20 mM Tris at pH 8.0, 500 mM NaCl), once with 1 mL LiCl wash buffer (0.25 M LiCl, 1% NP-40, 1% Na-deoxycholate, 1 mM EDTA, 10 mM Tris at pH 8.0) and twice with 1 mL 1x TE (10 mM Tris at pH 8.0, 1 mM EDTA). For each wash, the beads were rotated on a rotaspin for 5 min in the corresponding buffer, followed by centrifugation at 5400 rpm for 2 min. The beads were suspended in 0.25 mL of elution buffer (0.1 M NaHCO_3_, 1% SDS), incubated at 65 °C for 5 min, and rotated on the rotaspin for 15 min. The supernatant was collected after centrifugation. The elution step was repeated to obtain a final eluted volume of 0.5 mL. The elution buffer was also added to the stored input sample to obtain a final volume of 0.5 mL. Decrosslinking of the three samples (input, IP, and antibody-minus) was done by adding 20 μL of 5 M NaCl and overnight incubation at 65 °C. Proteins in the samples were removed by adding 10 μL 0.5 M EDTA, 20 μL 1 M Tris at pH 6.8, 2 μL Proteinase K (20 mg/L) and incubating at 45°C for 3 h. An equal volume of phenol-chloroform-isoamyl alcohol (25:24:1) was added for purifying the samples, and the aqueous phase was extracted by centrifugation at 14,000 rpm for 10 min. DNA was precipitated by adding 3 M Na-acetate (1/10th of the volume, pH 5.2), 1 μL glycogen (20 mg/mL), and 1 mL absolute ethanol, followed by incubation at −20°C overnight. The precipitated DNA was collected by centrifugation at 13,000 rpm for 30 min at 4°C and was washed once with 70% ethanol. Air-dried pellets were resuspended in 20 μL sterile MilliQ water with 10 mg/mL RNase. ChIP-DNA from duplicates were pooled for ChIP-seq.

The same protocol was followed to determine canonical histone H3 and histone H4 occupancy at the centromeres in *C. haemulonii*, *C. duobushaemulonii*, *C. pseudohaemulonii*, and different clades of *C. auris*, with some differences. Anti-H3 antibodies (Abcam [ab1791], at a final concentration of 13 μg/mL), and anti-H4 antibodies (Abcam [ab10158], at a final concentration of 13 μg/mL) were used for immunoprecipitation. The bead washes were done for 15 min.

### ChIP-seq

#### Library preparation

ChIP DNA obtained from CENP-A^Cse4^-Protein-A (4 ng) was used to generate a sequencing library using NEBNext^®^ Ultra^™^ II DNA Library Prep Kit for Illumina (Cat No. E7645S). In brief, the fragmented DNA was subjected to end repair followed by A – tailing and adapter ligation. The product DNA was enriched by PCR amplification using Illumina index adapter primers. The amplified product was purified using Ampure beads to remove unused primers. The library was quantitated using Qubit DNA High Sensitivity quantitation assay, and library quality was checked on Bioanalyzer 2100 using Agilent 7500 DNA Kit.

#### Data analysis

ChIP-sequencing yielded 20816547 reads for the input, and 20959149 reads for IP. Based on the FastQC (v0.11.8) report, adaptor sequences and orphan reads were removed using Trim Galore! (v0.4.4) (http://www.bioinformatics.babraham.ac.uk/projects/). The output file was mapped onto the GenBank reference assembly for *C. auris* clade 1 (GCA_002759435.2) to obtain the sequence alignment map in SAM format. Conversion to BAM, sorting, and indexing was achieved using SAMtools (v1.9)(Li et al. 2009). Identification and excision of duplicates were made using MarkDuplicates scripted by Picard tools (v1.119) (http://broadinstitute.github.io/picard/). The processed binary alignment map was used as input for MACS2 (v2.1.1) (Zhang et al. 2008) along with the genome control reads (processed in the same way as the immunoprecipitation sample) to generate peaks. The peaks were then sorted based on p-value, FDR value, and fold change. The peaks were visualized using Integrative Genomic Viewer (v2.4.1) (James T Robinson et al. 2011). Enrichment peaks were curated (fold enrichment ≥ 2.6), and the coordinates of the peaks obtained from MACS2 post-peak calling was used to extract sequences from the genome assemblies. The extracted sequences were scanned for repeats using SyMap (v4.2) (Soderlund et al. 2011), and the result was depicted as a dot plot.

### ChIP-qPCR analysis

Real-time PCR was used to confirm CENP-A^Cse4^ enrichment and H3 depletion in the centromere sequences, using primers specific to centromeres and non-centromeric loci (listed in supplementary table 3) and SensiFAST SYBR No ROX Kit. 1:50 dilutions for input and 1:20 dilutions of the IP were used for determining CENP-A ^Cse4^ enrichment. 1:50 dilutions for input and 1:5 dilutions of the IP were used for determining histone H3 and H4 occupancy. The program used: 94°C for 2 min, 94°C for 30 sec, appropriate T_m_ for 30 sec, 72°C for 30 sec for 30 cycles. The adjusted Ct values (log_2_ of dilution factor subtracted from the Ct value of the input or IP) were used to calculate the percentage input using the formula: 100*2^ (adjusted Ct of input-adjusted Ct of IP). Three technical replicates were taken for the assay, and the standard error of the mean was calculated. The plots were generated using GraphPad Prism 8.

### Ortholog search and Phylogenetic tree construction

Available annotation files for *S. cerevisiae* (GCF_000146045.2), *C. glabrata* (GCF_000002545.3), *C. albicans* (GCF_000182965.3), *C. tropicalis* (GCF_000006335.3), *C. dubliniensis* (GCF_000026945.1), *C. parapsilosis* (GCA_000182765.2), *D. hansenii* (GCF_000006445.2), *S. stipitis* (GCF_000209165.1), *C. neoformans* (GCF_000091045.1), *C. auris clade 1* (GCA_002759435.2), *C. auris clade 2* (GCA_003013715.2), *C. auris clade 4* (GCA_008275145.1), *C. duobushaemulonii* (GCF_002926085.2), *C. haemulonii* (GCF_002926055.2), *C. pseudohaemulonii* (GCF_003013735.1), *C. lusitaniae* (GCF_000003835.1), and *C. intermedia* (GCA_900106115.1) were downloaded from GenBank. Transcription and proteome data of *C. lusitaniae* were used to annotate the *C. fructus* (GCA_003707795.1) genome. *C. auris* clade 3 (GCA_005234155.1), *C. heveicola* (GCA_003708405.1), *C. oregonensis* (GCA_003707785.2), and *C. blattae* (GCA_003706955.2) genome assemblies were annotated using transcriptome and proteome data of *C. auris* clade 2, using MAKER (v2.31.10) (Holt & Yandell 2011). For all given species, clusters of orthologous proteins were identified using OrthoMCL (v2.0.9) (Li et al. 2003). The single-copy orthologs present in all the species were identified and aligned using Clustal Omega (v1.2.4) (Sievers et al. 2011). All the alignments were concatenated for each species, including the gaps. The gaps and corresponding sequences in all other species were removed. MrBayes (v2.3.5) (Ronquist & Huelsenbeck 2003) was used for tree construction, which was visualized using FigTree (v1.4.4) (http://tree.bio.ed.ac.uk/software/figtree/). Orthologs for proteins involved in heterochromatin formation and RNAi was done using phmmer option in HMMER (EMBL-EBI) (Potter et al. 2018).

### *In silico* analyses

#### Gene synteny

Centromere prediction in a candidate species was made by aligning the respective genome assembly to the reference species assembly using Mauve (Geneious v11.1.4) (Biomatters Ltd.), and the conserved synteny blocks corresponding to the ORFs flanking centromeres in the reference assembly were identified. For confirming synteny conservation, candidate species-specific local genome databases were created using Geneious. Blast analysis of five individual ORFs on either side of the centromeres in the reference species assembly was performed against the local genome database of the candidate species, using the protein sequences as queries. For genome-level comparison, coordinates of all the synteny blocks conserved between two species were obtained using SyMap (v4.2), and the circos plots were drawn using Circos (0.69-8) (Krzywinski et al. 2009). Scaffold-level and ORF-level synteny analyses identifying rearrangements were done using Easyfig (v2.2.2) (Sullivan et al. 2011).

#### Centromere sequence features

Python scripts were written to determine the GC% at the third position of codons. The percentage of G and C at the third position of codons (except the stop codons) was calculated, followed by calculating the average values in a sliding window of 10 ORFs. These values were plotted for each scaffold of the genome. Annotations that are not a multiple of three were not considered for the analysis. GC% was also calculated for the whole scaffolds with a window size of 5 kb and a sliding step of 1 kb. GC skew ((G – C)/ (G + C)) and AT skew ((A-T)/(A+T)) were plotted for a region of 10 kb flanking the centromeres using a window size of 100 bp and a sliding step of 1 bp. The skew calculation was done in Julia (v1.2.0), and the plotting was done in R. The “geom_smooth” function with “gam” method in ggplot2 (Wickham 2009) was used to smoothen the curve.

To study trends in centromere sequence evolution in different clades of *C. auris*, protein sequences were extracted using agat_sp_extract_sequences.pl from the AGAT suite (https://github.com/NBISweden/AGAT), and orthologous genes found using rsd_search (Wall et al. 2003). Intergenic sequence that occurred between the same pair of orthologous genes in pairs were identified as orthologous intergenic sequence and aligned using FSA (Bradley et al. 2009), which we previously found to have high specificity for true homology in aligning intergenic DNA sequence (Jayaraman & Siddharthan 2010). In each of the pairwise alignments generated by FSA, the mutation rate was estimated as #mutations/#matches, where #matches = number of positions where an aligned pair of nucleotides is reported; and #mutations = number of match positions where the alignment is a mismatch. The mean and sample standard deviation over all intergenic sequences were calculated and compared to the observed numbers in centromeres.

If available, the respective genome assembly annotation files were used to report the length of ORF-free regions. Otherwise, all predicted ORFs larger than 600 bp were considered as coding sequences. Motif search was done using MEME in the MEME Suite (Bailey et al. 2009).

#### Gene expression

For determining the transcriptional status of centromeres, the raw sequencing reads (SRR6900290, SRR6900291, SRR6900292, SRR6900293) were aligned to the reference genome of clade 1 (GenBank assembly GCA_002759435.2) using HISAT2 (v2.1.0) (Kim et al. 2019). The aligned reads were then graphically visualized in the IGV to analyse gene expression levels at/around the centromeres on different chromosomes. For studying the transcriptional status of ORFs overlapping with or flanking the centromeres, the abundance of annotated transcripts was quantified using pseudo alignment program kallisto (v0.46.1) (Bray et al. 2016). The expression of genes around/ overlapping the centromere in TPM (transcripts per million) were compared to the global gene expression level.

## Data availability

ChIP-sequencing data have been deposited in NCBI under BioProject PRJNA612018.

## Acknowledgements

We thank N. Chauhan for providing the strains, V. Yadav for preliminary gene synteny analyses, L. Sreekumar for reagents, and Clevergene Biocorp Pvt. Ltd., Bengaluru for generating ChIP-sequencing data. This study was funded by the Indian Council of Medical Research (AMR/149/2018-ECD-II), Government of India, to K.S., A.C., M.S., and R.P. K.S. acknowledges the financial support of JC Bose National Fellowship (Science and Engineering Research Board, Govt. of India, JCB/2020/000021) and intramural funding from JNCASR. A.N. was a National Postdoctoral Fellow (PDF/2016/003256), supported by the Science and Engineering Research Board (SERB), Department of Science and Technology (DST), Government of India. R.N.V, P.S, and R.S. acknowledge Department of Atomic Energy, Government of India for funding and the computing facilities provided by The Institute of Mathematical Sciences.

## Notes

### Competing Interest Statement

The authors have declared no competing interest.

