## Supplementary material for "Functional and comparative genome analysis reveals clade-specific genome innovations in the killer fungus *Candida auris*"

^1^ Molecular Mycology Lab, Molecular Biology and Genetics Unit, Jawaharlal Nehru Centre for Advanced Scientific Research, Bangalore, India; ^2^Computational Biology, The Institute of Mathematical Sciences/HBNI, Chennai, India; ^3^ [Postgraduate Institute of Medical Education and Research](https://scholar.google.co.in/citations?view_op=view_org&hl=en&org=8616802205652468316), Chandigarh, India; ^4^ Amity Institute of Biotechnology, Gurgaon (Manesar), Haryana, India.

Corresponding author

Kaustuv Sanyal

**Supplementary Fig. 1**

Degree of sequence conservation proving the identity of CENP-A ^Cse4^ in *C. auris* and related species. Amino acid sequences of CENP-A^Cse4^ from different ascomycetes were aligned using BioEdit (version 7.2). Key: *Scer*- *S. cerevisiae*, *Calb*- *C. albicans*, *Cdub*- *C. dubliniensis*, *Clus* – *C. lusitaniae*, *Cau* – *C. auris*, *Chae* – *C. haemulonii*, *C. duo*- *C. duobushaemulonii*, *Cphae* – *C. pseudohaemulonii*, *Cfruc*- *C. fructus***.** The diverging N-terminal tail is highlighted in grey and the conserved histone fold domain in green. The conserved structures within the histone fold domain are also shown.

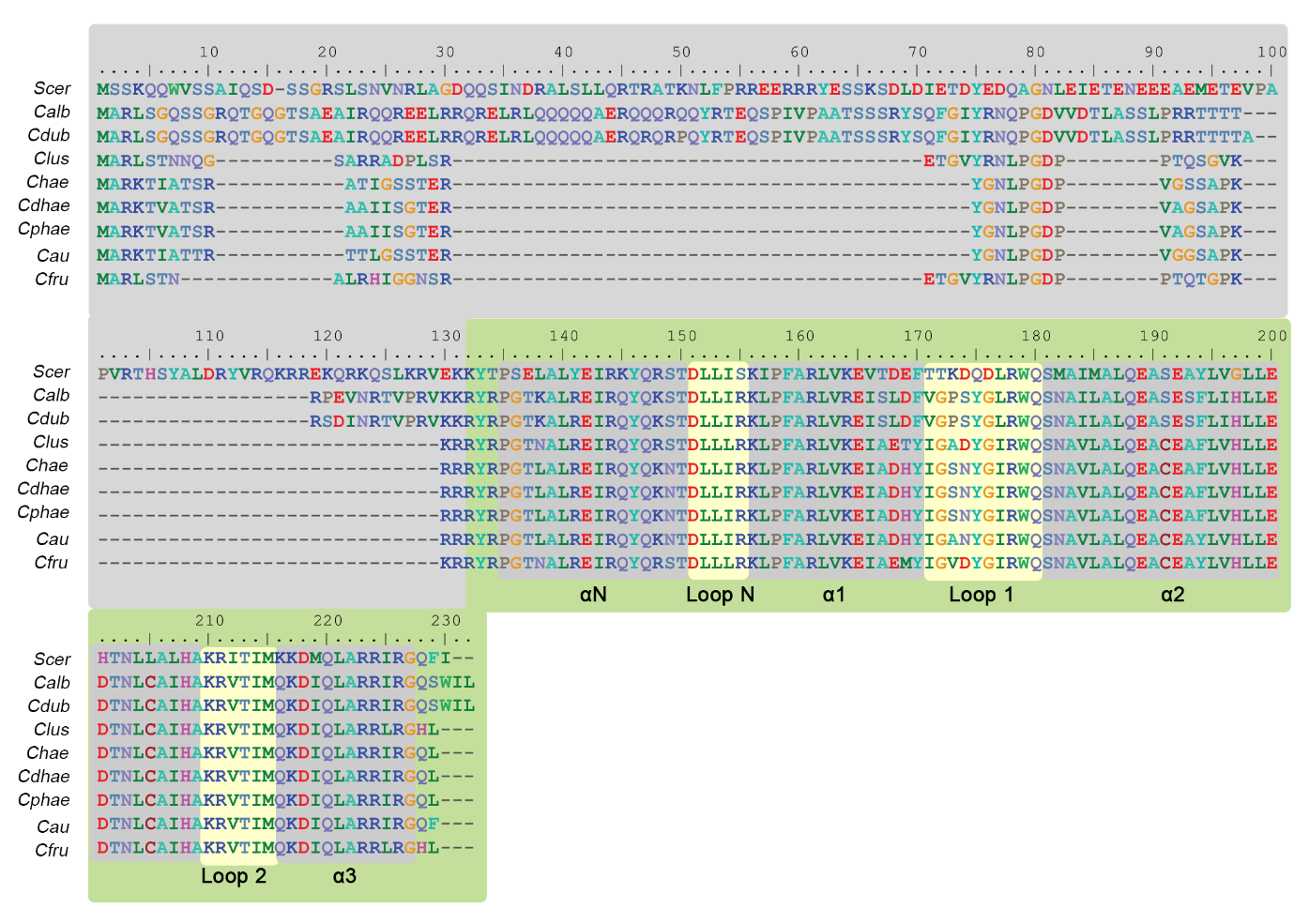

**Supplementary Fig. 2**

The AT-rich centromeres in *C. auris* lack pericentromeric heterochromatin. A, Design of tagging construct with the TAP tag and nourseothricin N-acetyl transferase (NAT) selection marker; western blot showing the expression of tagged CENP-A ^Cse4^. The size of the corresponding band in the ladder (middle lane) is marked. B, Dot plot comparing *CEN1* against itself identified the poly-A stretch (zoomed-in view, blue box). Motif search identified the poly-A stretch in all the centromere sequences (output logo from MEME suite, in red). C. Schematic comparing the lengths of the CENP-A ^Cse4^ enriched region and ORF-free region at the centromeres in *C. auris* (drawn to scale, shown on the x-axis kb). Box plots comparing the expression levels of D, 10 *CEN*-flanking genes, and E, *CEN*-overlapping genes with the global gene expression level. The middle yellow line shows the median value. The box represents the 25th percentile(Q1) to 75th percentile (Q3) values. The range of values(Q3-Q1) is the interquartile range (IQR). The whiskers represent Q1-(1.5*IQR) and Q3+(1.5*IQR). The remaining values which do not fall in the range are shown as outliers. F, Schematic showing the trend of disappearing ORF-free *CEN* neighbourhood in the *Clavispora/Candida* clade in Ascomycota. CENP-A ^Cse4^ enriched regions are indicated by dark grey bars and the ORF-free regions by the light grey bars. The nature of centromere in each case is also shown. The tree was drawn using PhyloT v2 (https://phylot.biobyte.de/).

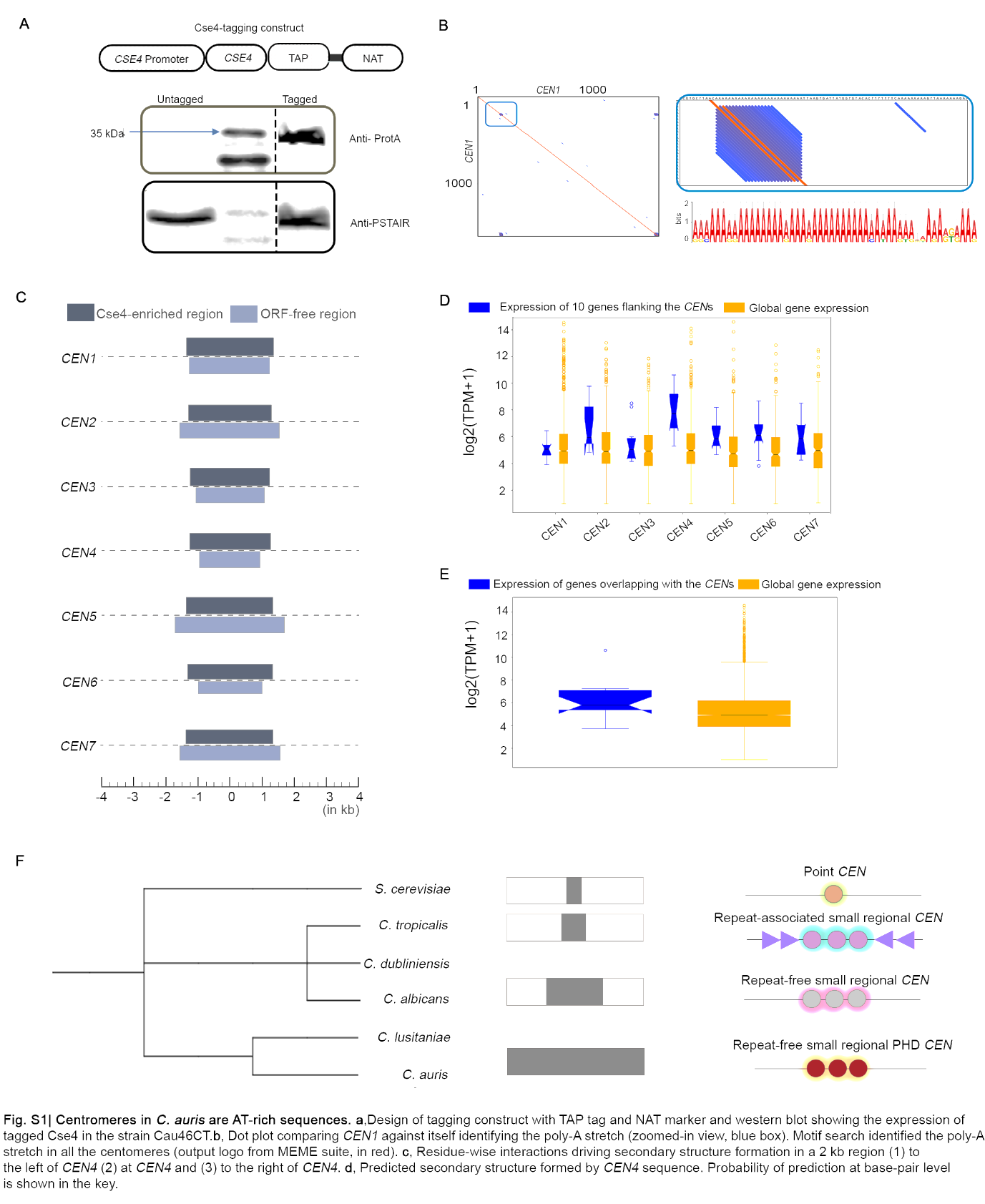

**Supplementary Fig. 3**

Centromere properties are conserved across the *C. auris* geographical clades. CENP-A ^Cse4^ enrichment at the *CEN*s in A, clade 2, D, clade 3, and G, clade 4. The corresponding depletion of canonical histone H3 in B, clade 2, E, clade 3, and H, clade 4 is depicted as H3/H4 ratio on the y-axis. The percent input values in all the experiments were compared to a control region (*far-CEN4*). qPCR values shown are from three technical replicates. The experiment was repeated twice, with similar results. Error bars indicate standard error of the mean (SEM). Statistical analysis was done using one-way ANOVA (**** P<0.0001, *** P<0.001). *CEN* positions (▲) overlap with GC- (▬) and GC-3 (▬) scaffold minima in C, clade 2 F, clade 3 I, clade 4. Coordinates (in Mb) are shown on the x-axis and %GC, on the y-axis.

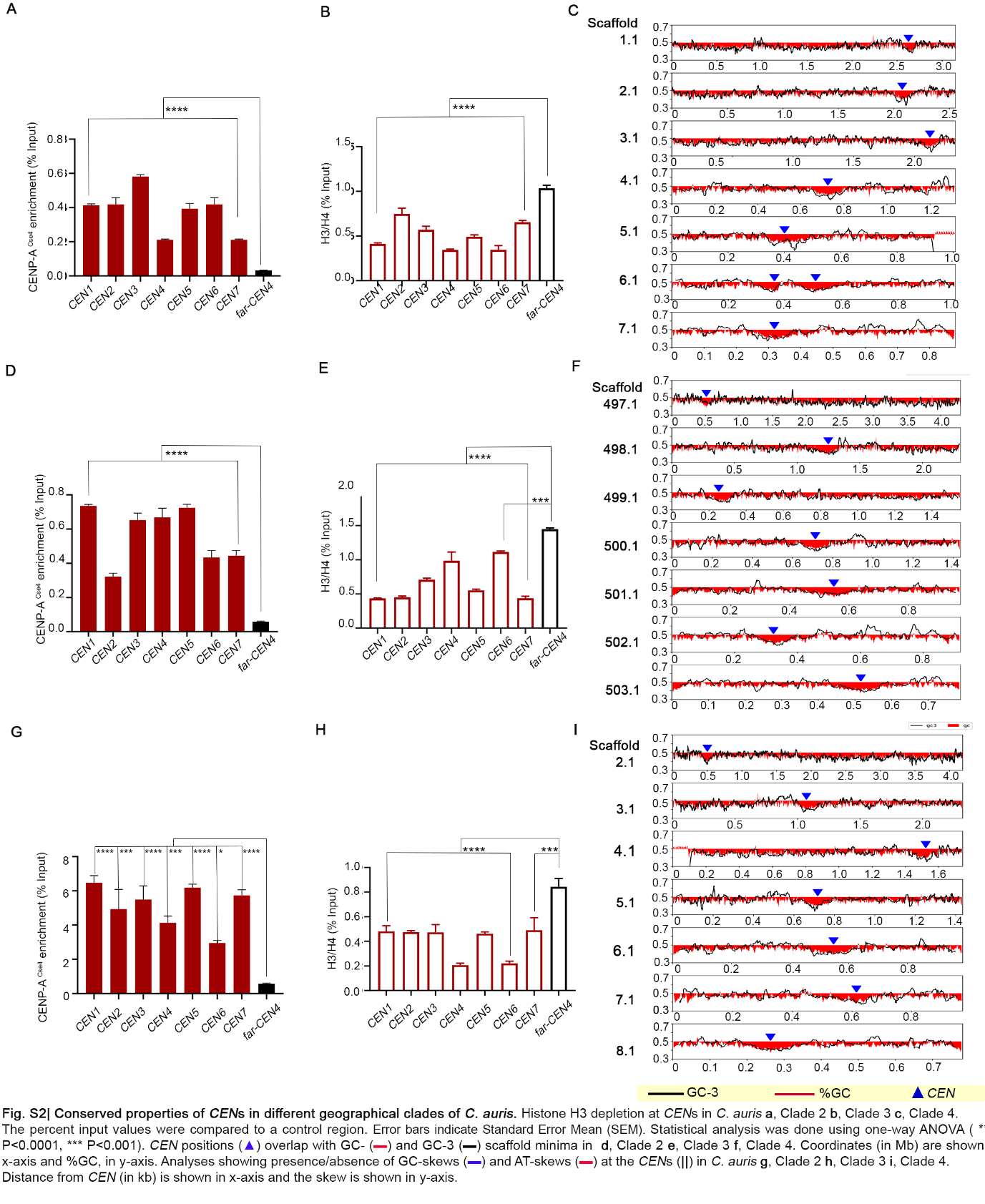

**Supplementary Fig. 4**

Putative centromeres in *C. haemulonii* and *C.pseudohaemulonii* have properties similar to *C. auris* centromeres. Circos plots showing synteny conservation between A, *C. auris* clade 4 and *C. haemulonii* e, *C. auris* clade 4 and *C. pseudohaemulonii*. Genomic scaffolds are shown on the outer-most track with the centromere positions marked by empty circles, the middle track shows %GC (red- GC content below genome average, blue- AT content above genome average), and the inner-most track depicts synteny blocks. *CEN* positions (▲) overlap with GC- (▬) and GC-3 (▬) scaffold minima in B, *C. haemulonii* F, *C. pseudohaemulonii.* Coordinates (in Mb) are shown on the x-axis and %GC, on the y-axis. Histone H3 depletion at the *CEN*s of C, *C. haemulonii* G, *C. pseudohaemulonii* is shown. The percent input values at *CEN*s on different scaffolds (shown on the x-axis) were compared to a non-centromeric control region (far-*CEN*). The values shown are from three technical replicates, and the experiment was repeated twice, with similar results. Error bars indicate standard error of the mean (SEM). Statistical analyses were done using one-way ANOVA, **** P<0.0001. Dot plot depicting the uniqueness of *CEN* sequences and absence of repeats in D, *C. haemulonii* H, *C. pseudohaemulonii*. Scaffold numbers are shown.

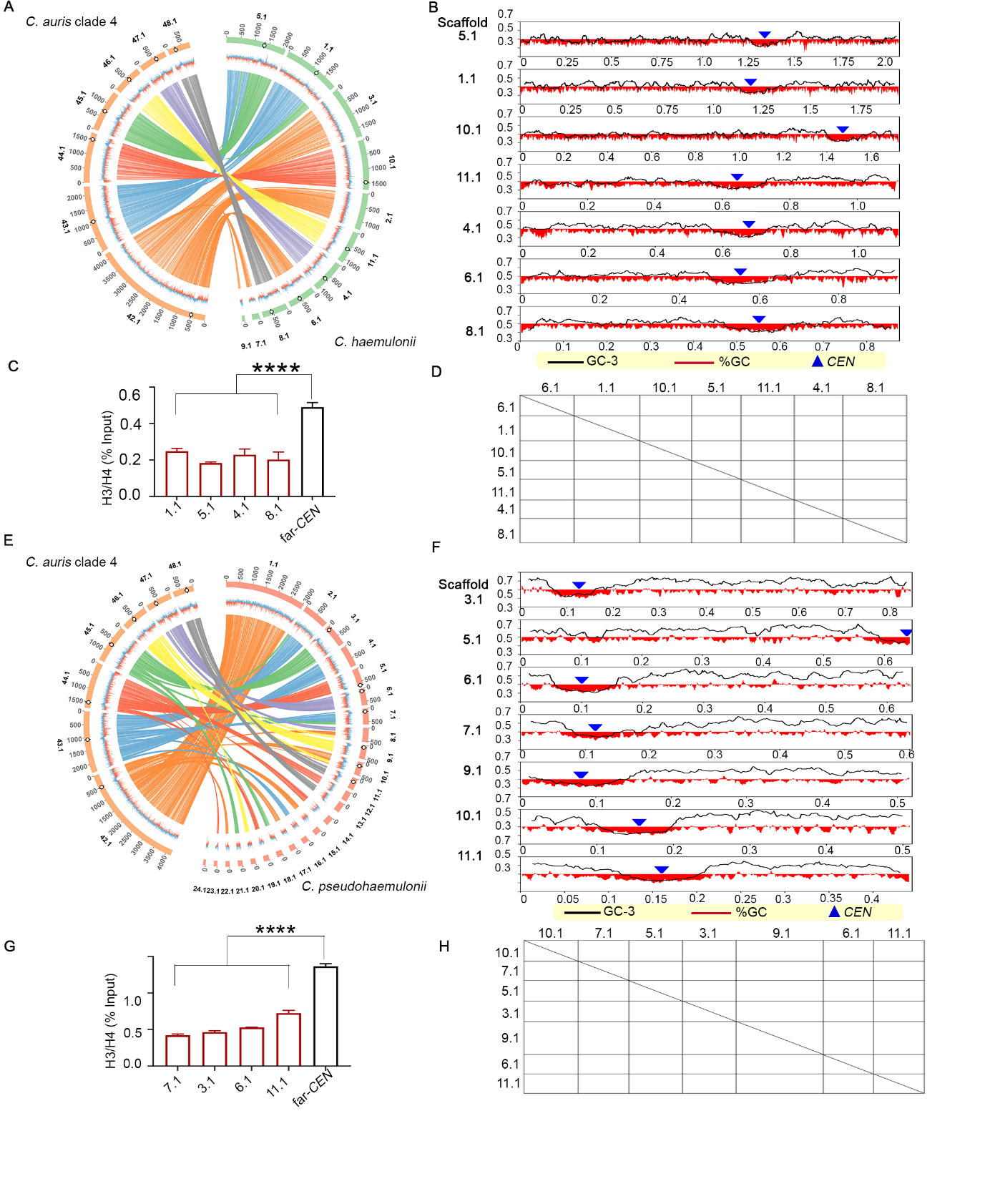

**Supplementary Fig. 5**

Genome-level comparisons reveal the degree of relatedness among species. Circos plot depicting chromosome-level similarities between A, *C. auris* clade 1 (strain CA-AM1) and *C. duobushaemulonii*, B, *C. auris* clade 3 (strain A1) and *C. duobushaemulonii* C, *C. auris* clade 2 and *C. duobushaemulonii*, D, *C. auris* clade 4 and *C. lusitaniae.* The outer-most track shows the genomic scaffolds with the *CEN* positions marked by empty circles, the middle track shows %GC (red- GC content below genome average, blue- AT content above genome average), and the inner-most track shows synteny blocks.

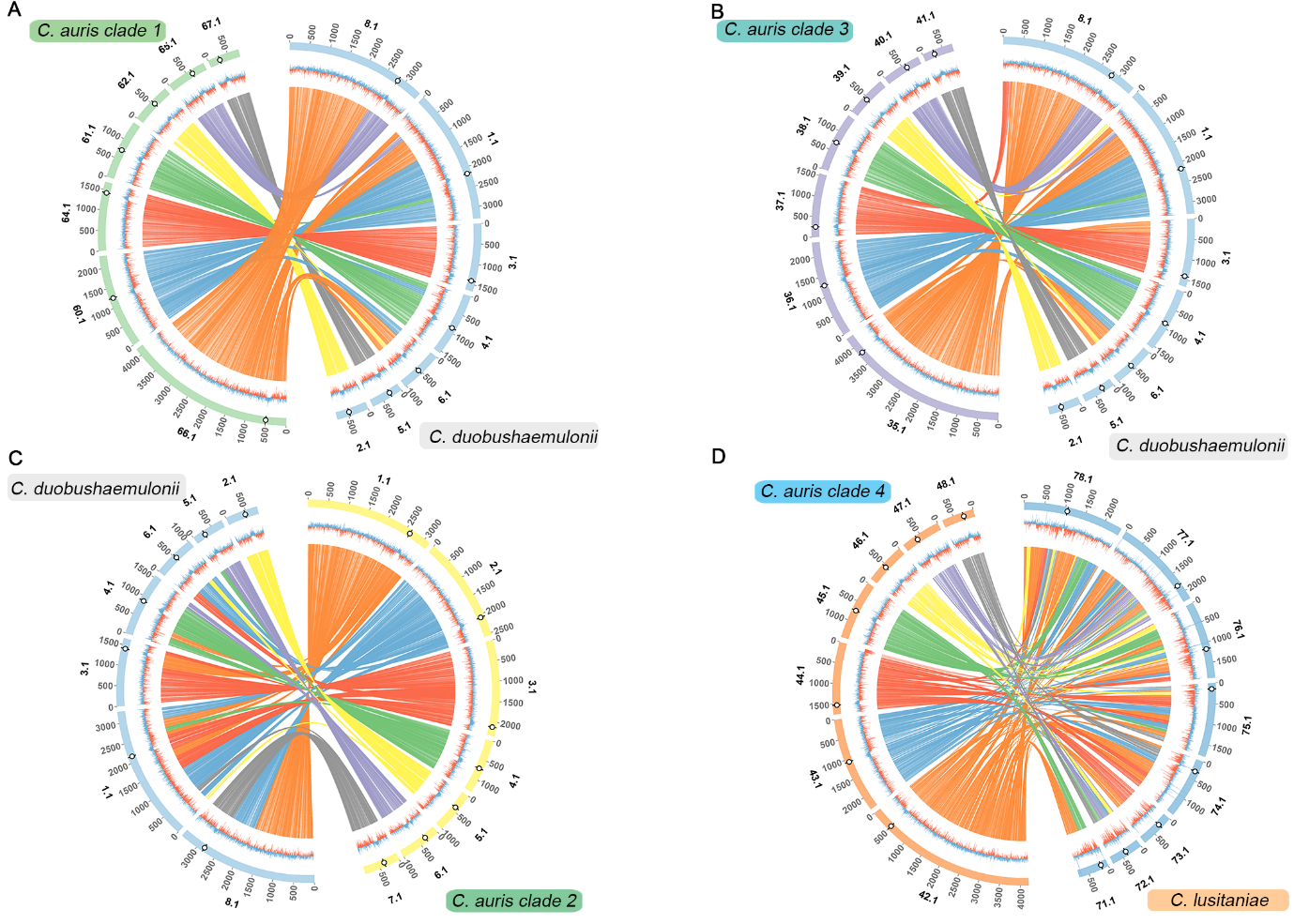

**Supplementary Fig. 6**

Centromere-associated structural changes in *C. haemulonii* and *C. pseudohaemulonii,* compared to *C. lusitaniae,* are similar to those in *C. auris*. Mapping of *C. lusitaniae CEN2* and *CEN8* synteny blocks onto the same scaffold in A, *C. haemulonii* B, *C. pseudohaemulonii*. Inactive *CEN* is shown as ▲. Pericentric inversion changing the relative position of ORFs 1,2 and 3 in C, *C. haemulonii* D, *C. pseudohaemulonii*. Separation of synteny blocks flanking *C. lusitaniae CEN3* by ~55 kb on the same chromosome in E, *C. haemulonii* F, *C. pseudohaemulonii*. Mapping of a synteny breakpoint at the centromere in G, *C. haemulonii,* and H, *C. pseudohaemulonii*. Centromeres in *C. lusitaniae* are marked by boxes (█ *C. lusitaniae*, █ *C. haemulonii*, and █ *C. pseudohaemulonii*). ▬ connects homologs, inversions (if present), are shown as ▬. The sequence similarity is shown as a percentage in the key.

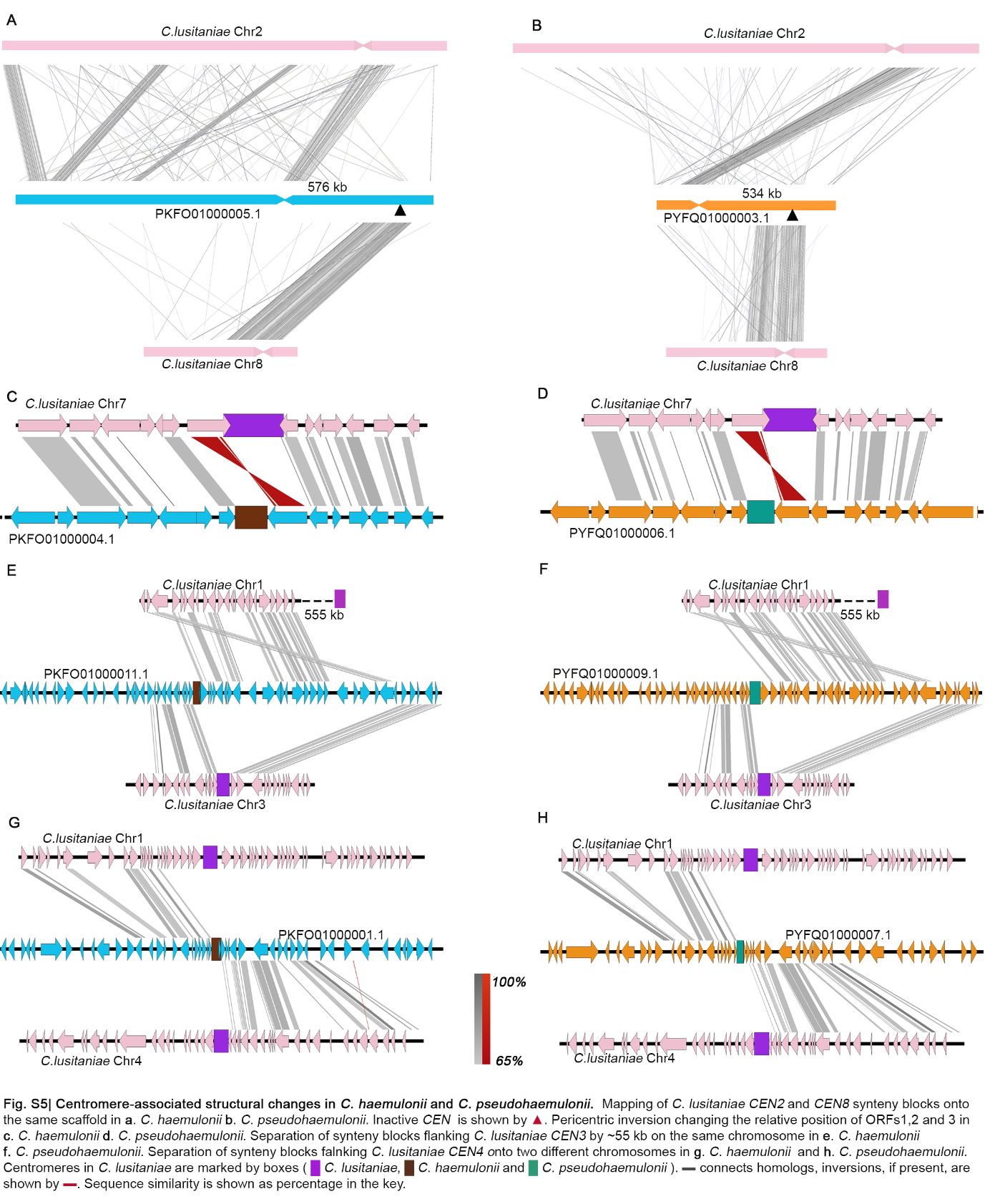

**Supplementary Fig. 7**

Eight putative centromeres identified in *C. fructus*. A, Eight loci in *C. fructus* syntenic to *C. lusitaniae* centromeres (█). Putative *CEN*s in *C. fructus* are marked by █. ▬ connects homologs, inversions (if present), are shown as ▬. The sequence similarity is shown as a percentage in the key. B, Putative *CEN* positions (▲) overlap with GC- (▬) and GC-3 (▬) scaffold minima. Coordinates (in Mb) are shown on the x-axis and %GC, on the y-axis. C, Plots showing the presence of GC-skews (▬) and AT-skews (▬) at the putative *CEN*s (||). Distance from *CEN* (in kb) is shown on the x-axis, and the skew is shown on the y-axis.

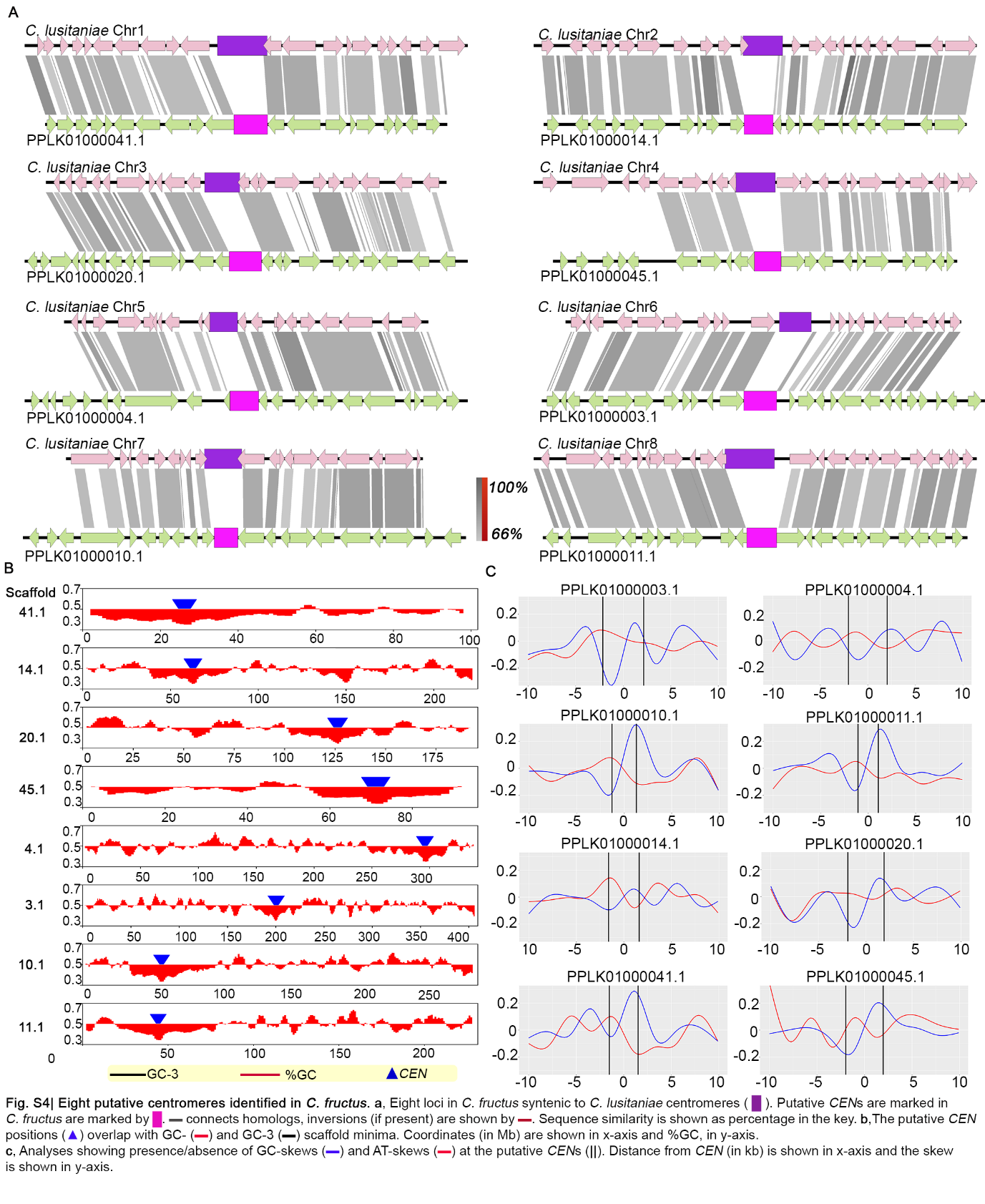

**Supplementary Fig. 8**

Conserved *CEN* neighbourhoods in other species of the *Clavispora/Candida* clade. A, Location of *CEN*s in *C. blattae*, *C. intermedia*, *C. heveicola,* and *C. oregonensis* were predicted based on gene synteny conservation, using *C. auris* as the reference. The colour code for ORFs and putative *CEN*s for each species is given in the key. ▬ indicates the GC-content, ▬ connects homologs, and inversions (if present) are shown as ▬. The sequence similarity is shown as a percentage in the key. B. Dot-plots showing the repeat-free, unique sequences at the centromeres in different species. Centromeres are numbered using *C. auris* as the reference. C. Plot comparing the lengths of centromeres identified/predicted in the study. Lengths of CENP-A ^Cse4^ enriched regions are shown for *C. lusitaniae* and *C. auris*. Lengths of the ORF-free region are shown for the rest of the species. The inactive *CEN* is depicted in black.

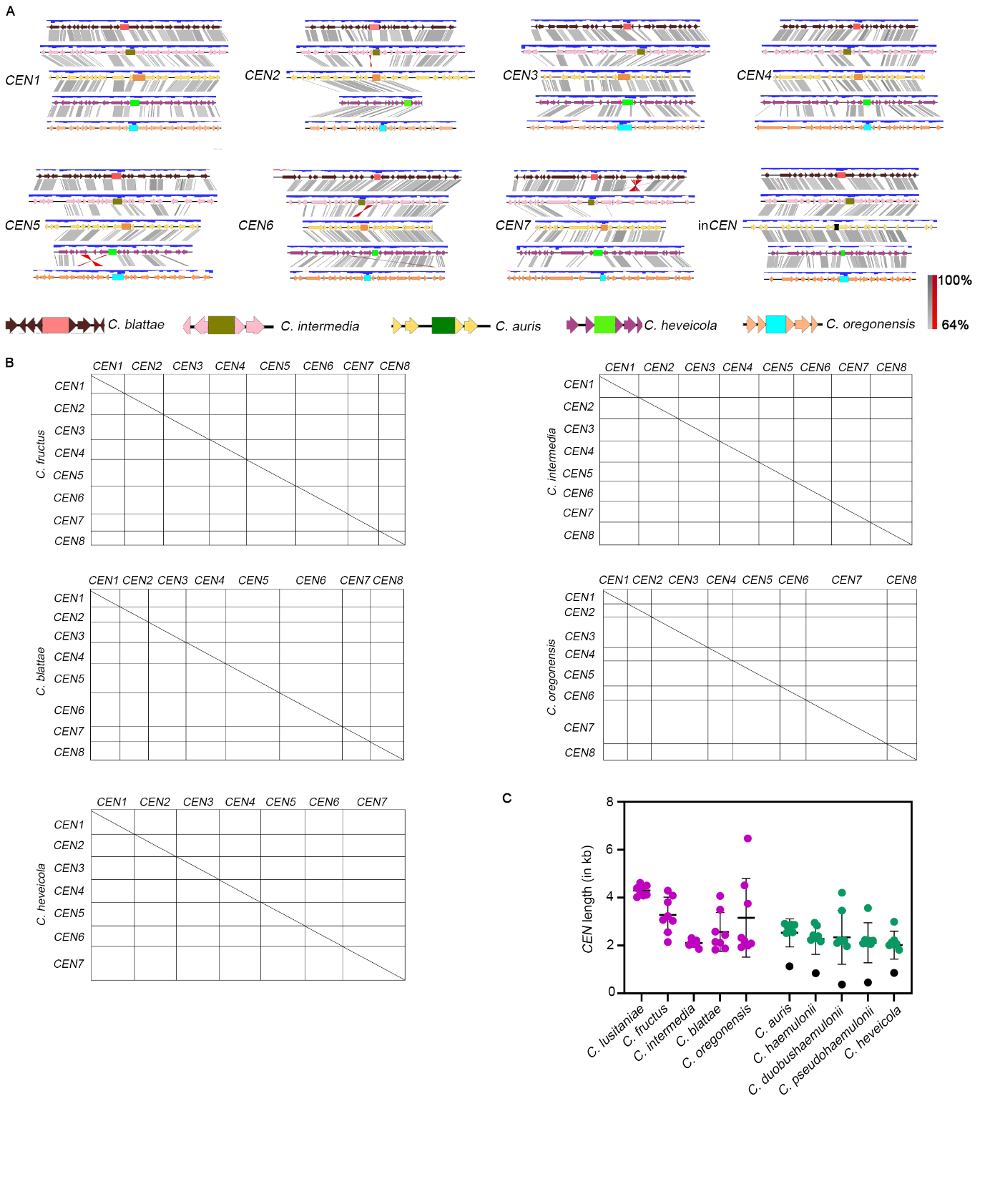

**Supplementary Table 1** Strains used in this study

| Strain | Species | Description | Source |
| --- | --- | --- | --- |
| Cau46R | *C. auris* | Clade 1 clinical isolate | NCCPF*, PGIMER, India |
| CBS1091131T | *C. auris* | Clade 2 strain | CBS |
| AR-0383 | *C. auris* | Clade 3 strain | Rutgers University |
| AR-0385 | *C. auris* | Clade 4 strain | Rutgers University |
| CauI46 | *C. auris* | *CSE4-TAP:: NAT* | This study |
| NCCPF470162 | *C. haemulonii* | Clinical isolate | NCCPF, PGIMER, India |
| NCCPF470163 | *C. pseudohaemulonii* | Clinical isolate | NCCPF, PGIMER, India |
| NCCPF470164 | *C. duobushaemulonii* | Clinical isolate | NCCPF, PGIMER, India |

*NCCPF- National Culture Collection of Pathogenic Fungi

**Supplementary Table 2** MLST analysis of *C. auris* clades

|  | *TUB2* | *RPB1* | *EFB1* |
| --- | --- | --- | --- |
| Base position* | 534 | 552 | 698 |
| Clade 1 | A | T | T |
| Clade 2 | A | C | T |
| Clade 3 | A | T | C |
| Clade 4 | G | T | C |

*with respect to clade 1

**Supplementary Table 3** Oligonucleotides used in this study

| Primer | Primer sequence (5’-3’) | Purpose |
| --- | --- | --- |
| D1-D2 seq FP | GCATATCAATAAGCGGAGGAAAAG | D1-D2 sequencing |
| D1-D2 seq RP | GGTCCGTGTTTCAAGACGG |  |
| TUB2 fp | AGATTACTCACTCCTTGGGT | MLST analysis |
| TUB2 rp | AGACAAGGTCGCGTTATATG |  |
| EFB1 fp | CTCAATGACCAAGTTGATCTG |  |
| EFB1 rp | AACTTGGATGAGTTGTTGGC |  |
| RPB1 fp | AAGAGCAAAATGGTCTGTGA |  |
| RPB1 rp | GCTCGTTTTCTTCAAACTGT |  |
| Au CEN1 fp | CCGATACAGTACATCGTTGA | ChIP-qPCR  *CEN* primers |
| Au CEN1 rp | TGATCAGTGGCGTTCAATAA |  |
| Au CEN1 fp | TGATAGACTTTTCTGGTGGGT | ChIP-qPCR  *CEN* primers  (clade 4) |
| Au CEN1 rp | TTCTACTTTATTACATGGCTTTGC |  |
| Au CEN2 fp | CCATTCGTTGCTTCAATCAT | ChIP-qPCR  *CEN* primers |
| Au CEN2 rp | TCGTCTATCGCTTCATACAC |  |
| Au CEN3 fp | CACGGTAATGGATTGACACTA | ChIP-qPCR  *CEN* primers |
| Au CEN3 rp | GTCAGGATTTCAGTGATGCT |  |
| Au CEN4 fp | TGATAGACTTTTCTGGTGGGT | ChIP-qPCR  *CEN* primers |
| Au CEN4 rp | TTCTACTTTATTACATGGCTTTGC |  |
| Au CEN5 fp | AACAGGCACACAGTCAGATG | ChIP-qPCR  *CEN* primers |
| Au CEN5 rp | CTGCGTAAGCTGAAAAACCG |  |
| Au CEN6 fp | ACACATTTACTTTTTGACGGG | ChIP-qPCR  *CEN* primers |
| Au CEN6 rp | GGGTAAGAATCGTGAGAGAA |  |
| Au CEN7 fp | GTAACCCATGTGGAGCACAA | ChIP-qPCR  *CEN* primers |
| Au CEN7 rp | CGGCCATGCATCAATCCTAT |  |
| Au CEN7 fp | TTCCATTCTCCAAGGATAGG | ChIP-qPCR  *CEN* primers  (clade 4) |
| Au CEN7 rp | TATCTAAATGCAATCGTGGG |  |
| Au 4532 fp | TATCTCTTGAGCTGGATGGT | ChIP-qPCR  Control region |
| Au 4532 rp | AACTTCCTGCTGGACAAAAT |  |
| Duo CEN1 fp | GGTCACTTATACGAACACCA | ChIP-qPCR  *CEN* primers |
| Duo CEN1 rp | TTACACGAGCTGCTATTACC |  |
| Duo CEN2 fp | CACAATCCTGTGATCTAGCA | ChIP-qPCR  *CEN* primers |
| Duo CEN2 rp | AATACGAAGCACTTCAACCT |  |
| Duo CEN3 fp | TTAACTGGTAATAGGGCACG | ChIP-qPCR  *CEN* primers |
| Duo CEN3 rp | AAAAAGTACGAAAACCGAGC |  |
| Duo CEN4 fp | ATGACCTAGGGACATCTTCT | ChIP-qPCR  *CEN* primers |
| Duo CEN4 rp | ATTTTGGAGACGACCATTTC |  |
| Duo 4532 fp | TAGATGTCAGGTGGTCAGAA | ChIP-qPCR  Control region |
| Duo 4532 rp | CGTTTTCATCAACACGCATA |  |
| Hae CEN2 fp | TTCTAAACAAACGTACCCGA | ChIP-qPCR  *CEN* primers |
| Hae CEN2 rp | AGCTAGGAGATGAAATCCGA |  |
| Hae CEN3 fp | ATGTAACGTCAATGGTGACA | ChIP-qPCR  *CEN* primers |
| Hae CEN3 rp | GCTCACAGTTTGTCTTAGGT |  |
| Hae CEN4 fp | TCTTAGTATGCCGTTGTAGC | ChIP-qPCR  *CEN* primers |
| Hae CEN4 rp | CGCCCAATCAAACACTAAC |  |
| Hae CEN6 fp | GCCTTTATGTCATGTTGTGC | ChIP-qPCR  *CEN* primers |
| Hae CEN6 rp | AACTATGTAGGCTGTTGCAT |  |
| Hae CEN7 fp | AGAGAAGTAGGTAGCTTGGA | ChIP-qPCR  *CEN* primers |
| Hae CEN7 rp | ATATGACCACATCAATCGGG |  |
| Hae 4532 fp | CCAGGAAGGTATTGAGGCTA | ChIP-qPCR  Control region |
| Hae 4532 rp | CCAACGTAGTCTACTCTTCTG |  |
| Phae CEN2 fp | CCTACTTCTGGCGTTCATAA | ChIP-qPCR  *CEN* primers |
| Phae CEN2 rp | AAAGCCTGCATTATTTGTCC |  |
| Phae CEN4 fp | CCATTCCCCTCCTTATTTGT | ChIP-qPCR  *CEN* primers |
| Phae CEN4 rp | GGAACTACAACCACCTGAAT |  |
| Phae CEN6 fp | ATTAGAAGCCTTGTTCGAGAT | ChIP-qPCR  *CEN* primers |
| Phae CEN6 rp | AACTAGAGACACCAACAGAG |  |
| Phae CEN7 fp | TTCTTAACAAGTGGTCGAGG | ChIP-qPCR  *CEN* primers |
| Phae CEN7 rp | AATCCGATCCACAACTTCAC |  |
| Phae 4532 fp | AACCACTACACATGGTCTTC | ChIP-qPCR  Control region |
| Phae 4532 fp | TCTTAGAAGCCTCCAAGTCT |  |
| CSE4-TAP-US-FP | AGCTGGTACCATGAGGATGGTGATTGAATCATC | Strain construction |
| CSE4-TAP-US-RP | agctACTAGTGAATTGACCTCGTATCCTTCG |  |
| CSE4-TAP-DS-FP | AGCTACTAGTATGAGAACTTTAGAGAATTGTGTC |  |
| CSE4-TAP-DS-RP | AGCTGCGGCCGCTCTGAGTCATAATATTCCGAGTC |  |
| CSE4-TAP- CN | GCTCAACAATACCGCAAATGTTATC |  |

**Supplementary Table 4** RNAi and heterochromatin-associated proteins in *C. auris* and *C. haemulonii* complex species

| Protein | Description | *C. auris* | *C. haemulonii* | *C. duobushaemulonii* | *C. pseudohaemulonii* |
| --- | --- | --- | --- | --- | --- |
| HP-1 | Chromodomain protein | **-** | **-** | **-** | **-** |
| Clr4 | Histone-lysine N-methyltransferase, H3 lysine-9 specific | **-** | **-** | **-** | **-** |
| Sir2 | NAD-dependent histone deacetylase SIR2 | B9J08_002917 | CXQ85_002270 | CXQ87_000715 | C7M61_000634 |
| Dcr1 | RNAse involved in RNA interference | B9J08_002318 | CXQ85_005187 | CXQ87_004766 | C7M61_003937 |
| Ago1 | RNA silencing | **-** | **-** | **-** | **-** |
| RdRP | RNA-dependent RNA polymerase | **-** | **-** | **-** | **-** |

**Supplementary Table 5** Centromeres in four geographical clades of *C. auris* – scaffold map

| *CEN* | Clade 1  Scaffold # | Clade 2  Scaffold # (Coordinates) | Clade 3  Scaffold # (Coordinates) | Clade 4  Scaffold # (Coordinates) | Strain CA-AM1 | Strain A1 |
| --- | --- | --- | --- | --- | --- | --- |
| *CEN1* | PEKT02000007.1 | CP043532.1:  (2073492-2075942) | CM016497.1:  (512473-514997) | CP043442.1:  (498292-500725) | CP061166.1, 506242-508839 | CP041135.1, 3784930- 3787568 |
| *CEN2* | PEKT02000001.1 | CP043533.1:  (2125357-2127677) | CM016498.1:  (1247622-1249916) | CP043443.1:  (1069183-1071379) | CP061160.1, 1241526- 1244388 | CP041136.1, 1281760- 1284636 |
| *CEN3* | PEKT02000003.1 | CP043531.1:  (2624933-2627082) | CM016499.1:  (251284-253638) | CP043444.1:  (1521159-1524233) | CP061164.1, 1432703- 1435251 | CP041137.1, 251504-254010 |
| *CEN4* | PEKT02000010.1 | CP043534.1:  (724286-726406) | CM016500.1:  (712095-714914) | CP043445.1:  (716366-718504) | CP061161.1, 735191-737700 | CP041138.1, 721942-724449 |
| *CEN5* | PEKT02000009.1 | CP043535.1:  (398873-401244) | CM016501.1:  (546843-550014) | CP043446.1:  (547070-549346) | CP061162.1, 550709-553616 | CP041139.1, 447449-450385 |
| *CEN6* | PEKT02000008.1 | CP043537.1:  (317145-319270) | CM016502.1:  (322203-324715) | CP043447.1:  (613673-615766) | CP061165.1, 627643-630375 | CP041140.1, 594422-597157 |
| *CEN7* | PEKT02000006.1 | CP043536.1**:**  (508176-510340)  (362335-364500) * | CM016503.1:  (515084-517108) | CP043448.1:  (262325-264369) | CP061167.1, 270648-273501 | CP041141.1, 271272-274040 |

* segmental duplication observed

**Supplementary Table 6** Centromere sequence evolution in different geographical clades

| Mutation rates | | | | | | | | | | | | |
| --- | --- | --- | --- | --- | --- | --- | --- | --- | --- | --- | --- | --- |
|  | **CENs (Z score)** | | | | **Inactive CENs (Z score)** | | | | **Intergenic (standard deviation)** | | | |
| Clade | 1 | 2 | 3 | 4 | 1 | 2 | 3 | 4 | 1 | 2 | 3 | 4 |
| 1 | 0 | 0.025 (0.273) | 0.092 (1.863) | 0.049 (0.583) | 0 | 0.011  (-0.131) | 0.082 (1.615) | 0.014  (-0.575) | 0 | 0.015 (0.034) | 0.013 (0.042) | 0.032 (0.030) |
| 2 | 0.025 (0.273) | 0 | 0.078 (5.119) | 0.051 (0.218) | 0.011  (-0.131) | 0 | 0.078 (5.125) | 0.020  (-0.497) | 0.015 (0.034) | 0 | 0.009 (0.013) | 0.042 (0.044) |
| 3 | 0.092 (1.863) | 0.078 (5.119) | 0 | 0.121 (5.106) | 0.082 (1.615) | 0.078 (5.125) | 0 | 0.089 (3.395) | 0.013 (0.042) | 0.009 (0.013) | 0 | 0.025 (0.019) |
| 4 | 0.049 (0.583) | 0.051 (0.218) | 0.121 (5.106) | 0 | 0.014  (-0.575) | 0.020  (-0.497) | 0.089 (3.395) | 0 | 0.032 (0.030) | 0.042 (0.044) | 0.025 (0.019) | 0 |

**Supplementary Table 7** Centromeres in the *C. haemulonii* complex species – scaffold map

| *C. auris CEN* | *C. haemulonii* | *C. duobushaemulonii* | *C. pseudohaemulonii* |
| --- | --- | --- | --- |
|  | Scaffold # (Coordinates) | Scaffold # (Coordinates) | Scaffold # (Coordinates) |
| *CEN1* | PKFO01000006.1  (551656-554089) | PKFP01000006.1  (533466-536672) | PYFQ01000010.1  (154148-156382) |
| *CEN2* | PKFO01000001.1  (1190716-1193669) | PKFP01000001.1  (2156789-2158893) | PYFQ01000007.1  (114370-116463) |
| *CEN3* | PKFO01000010.1  (1471204-1473502) | PKFP01000003.1  (1414893-1417085) | PYFQ01000005.1  (636137-638343) |
| *CEN4* | PKFO01000005.1  (1337308-1339535) | PKFP01000004.1  (885454-887675) | PYFQ01000003.1  (125155-127358) |
| *CEN5* | PKFO01000011.1  (642822-645213) | PKFP01000002.1  (465383-468840) | PYFQ01000009.1  (78470-82034) |
| *CEN6* | PKFO01000004.1  (676456-678623) | PKFP01000008.1  (2827810-2829780) | PYFQ01000006.1  (99229-101295) |
| *CEN7* | PKFO01000008.1  (549908-552739) | PKFP01000005.1  (275743-279947) | PYFQ01000011.1  (159135-161216) |

**Supplementary Table 8** Centromeres in *C. fructus* – scaffold map

| *C. lusitaniae CEN* | Scaffold # in *C. fructus* assembly | Coordinates |
| --- | --- | --- |
| *CEN1* | PPLK01000041.1 | 24682-27714 |
| *CEN2* | PPLK01000014.1 | 61157-64387 |
| *CEN3* | PPLK01000020.1 | 124314-128121 |
| *CEN4* | PPLK01000045.1 | 70020-73092 |
| *CEN5* | PPLK01000004.1 | 293788-297868 |
| *CEN6* | PPLK01000003.1 | 198330-202619 |
| *CEN7* | PPLK01000010.1 | 54115-56673 |
| *CEN8* | PPLK01000011.1 | 43113-45253 |

**Supplementary Table 9** Centromeres in *C. intermedia* – scaffold map

| *C. auris CEN* | Scaffold # in *C. intermedia* assembly | Coordinates |
| --- | --- | --- |
| *CEN1* | LT635762.1 | 97293-99600 |
| *CEN2* | LT635756.1 | 2116097-2118190 |
| *CEN3* | LT635761.1 | 421692-423715 |
| *CEN4* | LT635756.1 | 750835-752965 |
| *CEN5* | LT635758.1 | 677659-679836 |
| *CEN6* | LT635757.1 | 953334-955390 |
| *CEN7* | LT635759.1 | 597158-599012 |
| in*CEN* | LT635760.1 | 826270-828478*** |

*: No sequence loss

**Supplementary Table 10** Centromeres in *C. blattae* – scaffold map

| *C. auris CEN* | Scaffold # in *C. blattae* assembly | Coordinates |
| --- | --- | --- |
| *CEN1* | PPMS02000002.1 | 509108-511216 |
| *CEN2* | PPMS02000018.1 | 27373-29949 |
| *CEN3* | PPMS02000004.1 | 603681-607751 |
| *CEN4* | PPMS02000017.1 | 90553-92428 |
| *CEN5* | PPMS02000014.1 | 222093-224527 |
| *CEN6* | PPMS02000019.1 | 35853-37676 |
| *CEN7* | PPMS02000001.1 | 505461-508960 |
| in*CEN* | PPMS02000005.1 | 142874-145022* |

*: No sequence loss

**Supplementary Table 11** Centromeres in *C. heveicola* – scaffold map

| *C. auris CEN* | Scaffold # in *C. heveicola*  assembly | Coordinates |
| --- | --- | --- |
| *CEN1* | PPOB01000009.1 | 395236-397457 |
| *CEN2* | PPOB01000012.1 | 447091-449119 |
| *CEN3* | PPOB01000002.1 | 233988-236058 |
| *CEN4* | PPOB01000010.1 | 283258-285269 |
| *CEN5* | PPOB01000015.1 | 267746-269874 |
| *CEN6* | PPOB01000004.1 | 375189-377003 |
| *CEN7* | PPOB01000006.1 | 552389-555379 |

**Supplementary Table 12** Centromeres in *C. oregonensis*– scaffold map

| *C. auris CEN* | Scaffold # in *C. oregonenesis*  assembly | Coordinates |
| --- | --- | --- |
| *CEN1* | PPLJ02000002.1 | 391512-393700 |
| *CEN2* | PPLJ02000010.1 | 186066-187999 |
| *CEN3* | PPLJ02000003.1 | 203021-207534 |
| *CEN4* | PPLJ02000005.1 | 151694-153692 |
| *CEN5* | PPLJ02000008.1 | 277804-281557 |
| *CEN6* | PPLJ02000001.1 | 1139147-1141232 |
| *CEN7* | PPLJ02000007.1 | 451609-458079 |
| in*CEN* | PPLJ02000013.1 | 162814-165140* |

*: No sequence loss

**Supplementary Table 13** Distribution of *C. lusitaniae* *CEN8*-containing synteny block in *C. auris* clades and related species

| Synteny block in *C. lusitaniae*  Scaffold (start-end) | Size (in bp) | Synteny block in | Scaffold (start-end) | Size (in bp) |
| --- | --- | --- | --- | --- |
| CH408083.1  (89697-359662) | 269965 | *C. auris* clade 1 | PEKT02000010.1 (1066463-1295990) | 229527 |
| CH408083.1  (89707-217179) | 127472 | *C. auris* clade 2 | CP043536.1 (127217-235434) | 108217 |
| CH408083.1  (243462-359663) | 116201 | *C. auris* clade 2 | CP043534.1 (1076434-1178122) | 101688 |
| CH408083.1  (89707-359645) | 269938 | *C. auris* clade 3 | CM016500.1 (1065258-1294427) | 229169 |
| CH408083.1  (89707-359663) | 269956 | *C. auris* clade 4 | CP043445.1  (134688-363058) | 228370 |
| CH408083.1  (89745-359649) | 269904 | *C. haemulonii* | PKFO01000005.1  (1736412-1973270) | 236858 |
| CH408083.1  (89640-359720) | 270080 | *C. duobushaemulonii* | PKFP01000004.1  (1235993-1464114) | 228121 |
| CH408083.1  (89719-356554) | 266835 | *C. pseudohaemulonii* | PYFQ01000003.1  (485191-714677) | 229486 |

**Supplementary Table 14** *CEN*-associated changes in different species

| Species | SB (*CEN*3) | SB(*CEN*4) | PI | in*CEN* |
| --- | --- | --- | --- | --- |
| *C. lusitaniae* |  |  |  |  |
| *C. fructus* |  |  |  |  |
| *C. intermedia* |  |  |  |  |
| *C. blattae* |  |  |  |  |
| *C. oregonensis* |  |  |  |  |
| *C. auris* |  |  |  |  |
| *C. haemulonii* |  |  |  |  |
| *C. duobushaemulonii* |  |  |  |  |
| *C. pseudohaemulonii* |  |  |  |  |
| *C. heveicola* |  |  |  |  |

SB- Synteny break (*CEN*# with respect to *C. lusitaniae CEN*), PI-Pericentric Inversion, in*CEN*- Inactive *CEN* (assessed by sequence loss). Presence of the event is shown in black and the absence, in grey.
